# The carnosinase dipeptidase CNDP1 is a novel metabolic vulnerability in brain metastasis

**DOI:** 10.1101/2025.03.18.644053

**Authors:** Maria A. Gomez-Munoz, Alcida Karz, Maya Navarro, Victoria Osorio-Vasquez, Olga Katsara, Adam Walker, Pietro Berico, A. Paulina Medellin, Nicole M. Eskow, Yunjing Wan, Milad Ibrahim, Somnath Tagore, Edridge K. D’Souza, Benjamin Izar, Eleazar Vega Saenz de Miera, Iman Osman, Michael Pacold, Xiangpeng Kong, Mitchell Levesque, Richard Possemato, Sheri L. Holmen, Robert J. Schneider, C. Theresa Vincent, Kelly V. Ruggles, Douglas E. Biancur, Eva Hernando

## Abstract

Brain metastatic cells undergo metabolic adaptations, such as increased reliance on oxidative phosphorylation. Integrating proteomic and transcriptomic profiling of patient samples revealed a consistent upregulation of the carnosine dipeptidase-1 (CNDP1) in brain versus extracranial metastases. Carnosine is an abundant metabolite in brain and muscle, known to exert anti-proliferative effects on cancer cells. Here, we demonstrate that CNDP1 inhibition suppresses both the establishment and maintenance of melanoma brain metastasis while its ectopic expression is sufficient to confer brain metastatic potential to poorly metastatic cells. CNDP1 suppression results in activation of the Integrated Stress Response via Heme-Regulated Inhibitor Kinase and reprogrammed translation towards preferential expression of mitochondrial and survival transcripts. We further show that CNDP1 upregulation supports mitochondrial activity by limiting the levels of its substrate carnosine, a copper ionophore, thus protecting metastatic cells from carnosine-induced copper toxicity. Our studies reveal a novel metabolic adaptation during brain metastasis, which can be leveraged for therapeutic purposes.

## Introduction

While advances in cancer therapy have drastically improved patient care, metastatic disease is often refractory to current treatment options and remains the most common cause of cancer-related deaths(1,2). Primary tumor cells may acquire mutations or transcriptional programs that predispose cells to disseminate(3–5), and once they reach the metastatic niche, they must undergo further site-specific adaptations to survive in the new microenvironment (TME)(6–8). Understanding these adaptations is key to uncovering novel and targetable vulnerabilities of metastatic cells.

Specifically, brain metastases (BMs) are a major cause of cancer-related morbidity and mortality, affecting nearly 40% of patients with advanced solid tumors, most frequently in patients with carcinomas of the lung or breast (9–11), and melanoma(12). Despite tremendous advances in surgical approaches, radiotherapy, and systemic targeted and immune therapies, patients with symptomatic BMs experience reduced treatment response rates, even when extracranial metastases (ECMs) are controlled(13–15). Therefore, there is an urgent need of improved treatment options tailored to the unique BM biology(16–20).

Extensive efforts have been made to identify mechanisms of brain metastasis tropism (21–25) . For instance, studies have shown that metastatic cells rewire their metabolism to adapt to the distinctive properties of the brain such as higher oxygen concentration, distinct micronutrient availability, amino acid deprivation or alternative sources of carbon(22,23,26,27). Genes involved in oxidative phosphorylation have been found to be upregulated at both mRNA and protein levels in melanoma brain metastasis (MBM) patient samples compared to extracranial metastases(22,26), and patients with an increased oxidative phosphorylation signature in their MBMs exhibit decreased survival(21,26). We have previously shown that melanoma cells derived from the brain metastatic niche hold a higher oxidative potential than their extracranial counterparts(25). Moreover, metastatic cells undergo metabolic adaptations that enhance their capacity to survive under oxidative stress(28,29), such as modulating the expression of NADPH+ generating enzymes. Similarly, cells that metastasize to the leptomeninges are able to compete for available iron with macrophages in the cerebrospinal fluid (CSF) by modulating lipocalin-2 expression in the hypoxic niche to support their metabolic needs, among other cellular processes(30). A better understanding of metabolic dependencies in the brain metastatic niche may reveal novel susceptibilities that can be exploited for therapeutic purposes.

Here, we integrated two datasets: mass spectrometry-based proteomics of melanoma extracranial (non-brain metastasis; NBM) and MBM Short Term Cultures (STCs)(25), and transcriptomics data from extracranial and brain metastasis samples(26), reasoning that gene products consistently upregulated in MBM across different cohorts and two omics modalities might have functional relevance to the MBM cascade. This integration revealed that expression of the brain-associated dipeptidase CNDP1 is consistently increased in MBM versus NBM. CNDP1, or carnosine dipeptidase-1, is a secreted peptidase known to cleave carnosine into its component amino acids L-histidine and ß-alanine(31–34). Carnosine exerts a protective physiological function in muscle, brain, liver and kidney, and is reported to have anti-glycation effects and impact oxidative stress, though the respective underlying mechanisms, as well as those governing its abundance, remain unclear(35–38). It has been recently suggested that these properties of carnosine come from its ability to chelate metals such as copper and iron, which can contribute to the formation of Advanced Glycation End (AGE) products through reactive carbonyl or reactive oxygen species (ROS)(37–42). Carnosine administration *in vitro* is known to exert an anti-proliferative effect on cell lines of various cancer types(34,43–45). We therefore hypothesized that CNDP1 upregulation in MBM may provide an adaptative advantage to cancer cells in the brain where abundant carnosine exhibits a tissue-protective role. Indeed, our work uncovers a new role for CNDP1 in overcoming metabolic stress during MBM establishment by cleaving carnosine excess and protecting cells from intracellular copper overload. Our studies reveal a novel molecular mechanism of BM adaptation, and CNDP1 as a druggable vulnerability which can be exploited therapeutically to prevent or treat BM.

## Results

### Integrative multi-omics approach nominates CNDP1 as a driver of MBM

To uncover mechanisms of adaptation to the brain microenvironment shared across patients, proteomics data from paired NBM and MBM short-term cultures (STCs) (n=3 unique patients) were compared to an RNA-sequencing (RNA-seq) dataset of 35 surgically resected MBM and 41 NBM tissue specimens (n=29 unique patients, Table S1(25,26)). We compared differentially expressed genes in paired MBM vs NBM samples to those found by proteomics profiling (Figure 1A). By applying a cutoff of log2FC > 0.4 in both studies, and a p-value threshold of 0.005 in the differential transcriptomic analysis, we compiled a list of candidate genes upregulated in both cohorts, with Carnosine dipeptidase-1 (CNDP1) having the highest upregulation on both datasets (Figure 1A, Table S2). CNDP1 is a secreted zinc metalloprotease that cleaves the dipeptide carnosine into L-histidine and ß-alanine(46). *CNDP1* is predominantly expressed in the mammalian brain and liver, while its homologue *CNDP2* and its counterpart in carnosine metabolism *CARNS1* (carnosine synthase 1) are ubiquitously expressed (Figure S1A). Data mining of SKCM TCGA (The Cancer Genome Atlas) transcriptomics data revealed that *CNDP1* expression is significantly increased in metastatic (n=367) compared to primary samples (n=104) (p=0.0054; Figure S1B). *CARNS1* levels were found increased and *CNDP2* levels decreased during progression from primary to metastasis whereas CNDP1 increased in metastatic (n=367) compared to primary samples (n=104) (p<0.006, p<0.001 and p<0.06, respectively; Figure S1B). Neither *CNDP1* nor *CNDP2* transcript levels associate with melanoma patients’ overall survival in that dataset, while lower *CARNS1* correlates with shorter overall survival in metastatic patients (p<0.002; Figure S1C). In a separate cohort, CNDP1 protein levels within melanoma-infiltrated lymph nodes associate with shorter patient survival (n=59, p=0.03, Figure 1B). Additionally, RNA-sequencing analysis of melanoma extended to resected lymph nodes revealed higher expression of *CNDP1* in melanomas that metastasized to brain relative to those that recurred to other sites (p=0.03, Figure 1C, Table S4). Single nucleus RNA-sequencing (snRNA-seq) data from fresh-frozen patient samples(47) showed a higher percentage of MBM cells expressing *CNDP1* compared to NBM (Figure 1D; p = 6.22e^−16^). Extending our findings beyond melanoma, analysis of lung and breast cancer transcriptomics datasets showed *CNDP1* upregulation in BM samples relative to primary tissues (p<0.001; Figure 1E-F, S1D). Immunohistochemistry analysis of a tissue microarray (TMA) of melanoma patient samples confirmed higher CNDP1 levels in brain metastasis samples (n=70) compared to patient-matched NBM (n=24), primary tumors (n=43) or healthy skin and cortex (p=0.02, Figure 1G, Table S3). We confirmed the expression of CNDP1 by qPCR, immunoblotting and carnosinase activity in patient-matched STCs (Figure S1D-F). Altogether, these data nominated CNDP1 as a potential driver of brain metastasis.

**Figure 1.**
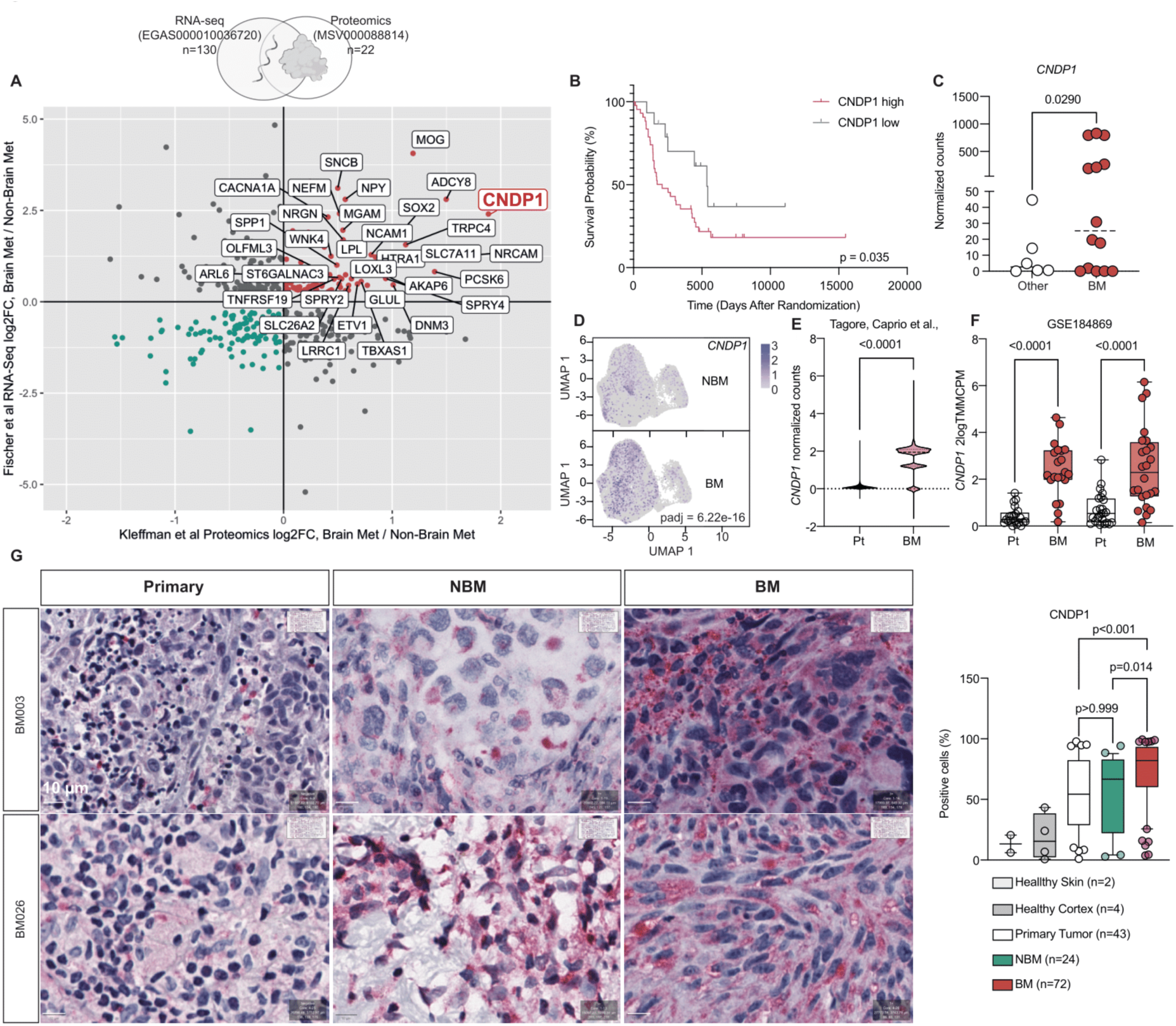
Multi-omics approach nominates *CNDP1* as driver of melanoma brain metastasis. **(A)** Data integration strategy showing log2Fold expression changes between patient-matched Brain Metastasis (BM) and Non-Brain Metastasis (NBM) tissue samples (RNA-sequencing n = 29 patients with the following samples: 35 brain, 7 intestine, 18 lymph node, 5 lung, 12 skin. Proteomics, n = 3 patients with the following samples: 3 brain, 1 subcutaneous, 1 bowel). NBM sources in RNA-sequencing data included P value (edgeR) < 0.005 for differential expression in RNA-sequencing (EGAS000010036720, Fischer et al.) No p-value threshold was applied for proteomics data. Genes labeled are log2FC > 0.4 in both studies. Additional data can be found in Table S1 and S2. **(B)** Kaplan-Meier curve showing patient Overall Survival stratified by high and low CNDP1 protein levels in lymph nodes metastasis on MM500 study(130). Cut off established between the lower and upper quartiles of protein intensity values. T test by log-rank test. Corrected for 50% minimum tumor content. **(C)** *CNDP1* differential mRNA expression represented as normalized counts in melanoma samples spread to the lymph node that later on metastasize to the brain or to other organs. Statistical analysis by Well’s t-test. Additional data can be found in Table S4. **(D)** Single Nucleus RNA-sequencing (snRNA-seq) data of human patient samples mined from Biermann et al (Series GSE185386). UMAP representing *CNDP1* expression in extracranial metastasis (NBM) vs intracranial metastasis (MBM). Statistics derived from Seurat’s FindMarkers function which applies a Wilcoxon Rank Sum test. **(E)** *CNDP1* normalized counts of single cell RNA-seq data of malignant cells (n=333050 cells) from lung adenocarcinoma primary tumors (LPT) and lung adenocarcinoma brain metastasis (LBM). Data from Tagore, Caprio et al. *in press* in Nature Medicine. Statistical analysis by Welch’s test. **(F)** *CNDP1* differential mRNA expression (2log TMMCPM) in breast cancer (BC) versus breast cancer brain metastasis (BCBM) paired patients’ samples from indicated data set. Statistical analysis by paired t-test. **(G)** Immunohistochemistry (IHC) showing differential expression of CNDP1 in patient-matched primary, extracranial metastasis and brain metastatic samples (TMA, n samples indicated). **Left.** Representative primary, extracranial and brain metastasis IHC DAB images (BM003, BM0026) Scale=10uM**. Right.** CNDP1 expression quantification in healthy skin, healthy cortex, primary tumor, NBM (no brain metastasis) and BM (brain metastasis) samples represented as % of positive cells. P value by Paired One way ANOVA. Additional data included in Table S3. See Figure S1

Next, we investigated potential transcriptional programs driving *CNDP1* upregulation in melanoma metastasis. *In silico* analysis of TCGA ATAC-seq data revealed open chromatin regions in the *CNDP1* locus and identified an enhancer, the opening of which strongly correlates with *CNDP1* expression (Fig. S1H-I). It has been reported that levels of *MITF*, the master regulator of melanocyte expression (48), are significantly higher in BM versus NBM (47), suggesting *CNDP1* transcription could be driven by MITF. ChIPseq confirmed direct MITF binding to the *CNDP1* promoter and enhancer (Fig. S1J), and MITF silencing resulted in decreased H3K27ac at the *CNDP1* promoter and reduced *CNDP1* mRNA levels (Figure S1J-L). Together, our data suggests that *CNDP1* expression may be at least partially governed by MITF.

### CNDP1 depletion and carnosine accumulation decrease *in vitro* proliferation of MBM patient-derived short-term cultures

To test the potential role of CNDP1 in MBM, we used a loss-of-function approach. CNDP1 silencing with two doxycycline (dox)-inducible shRNAs in two human MBM STCs (Figure 2A) slowed cell proliferation after 24 h by G1 phase stalling (p<0.001, Figure 2B-C), suggesting that CNDP1 is required for 2D cell growth in MBM cells. Importantly, 24 hours of dox-induced CNDP1 KD resulted in an increase in carnosine extracellular levels measured by ELISA (Figure S2A). We then confirmed that carnosine supplementation at comparable concentrations to those found physiologically in brain (49), leads to a proliferation defect in the same cells and increased intracellular carnosine levels (p<0.001, Figure S2B-C). Additionally, CNDP1 dependency is not a general feature of melanoma cells, as suggested by a proliferation assay in a non-brain tropic melanoma cell line (Figure S2D). We confirmed the anti-proliferative effect of CNDP1 depletion using RNA interference with pooled siRNA as an orthogonal approach (p<0.001; Figure 2D-E). Taken together, these data suggest cellular reprogramming to the brain microenvironment involves CNDP1 induction, exposing a novel and potentially targetable vulnerability of brain metastases.

**Figure 2.**
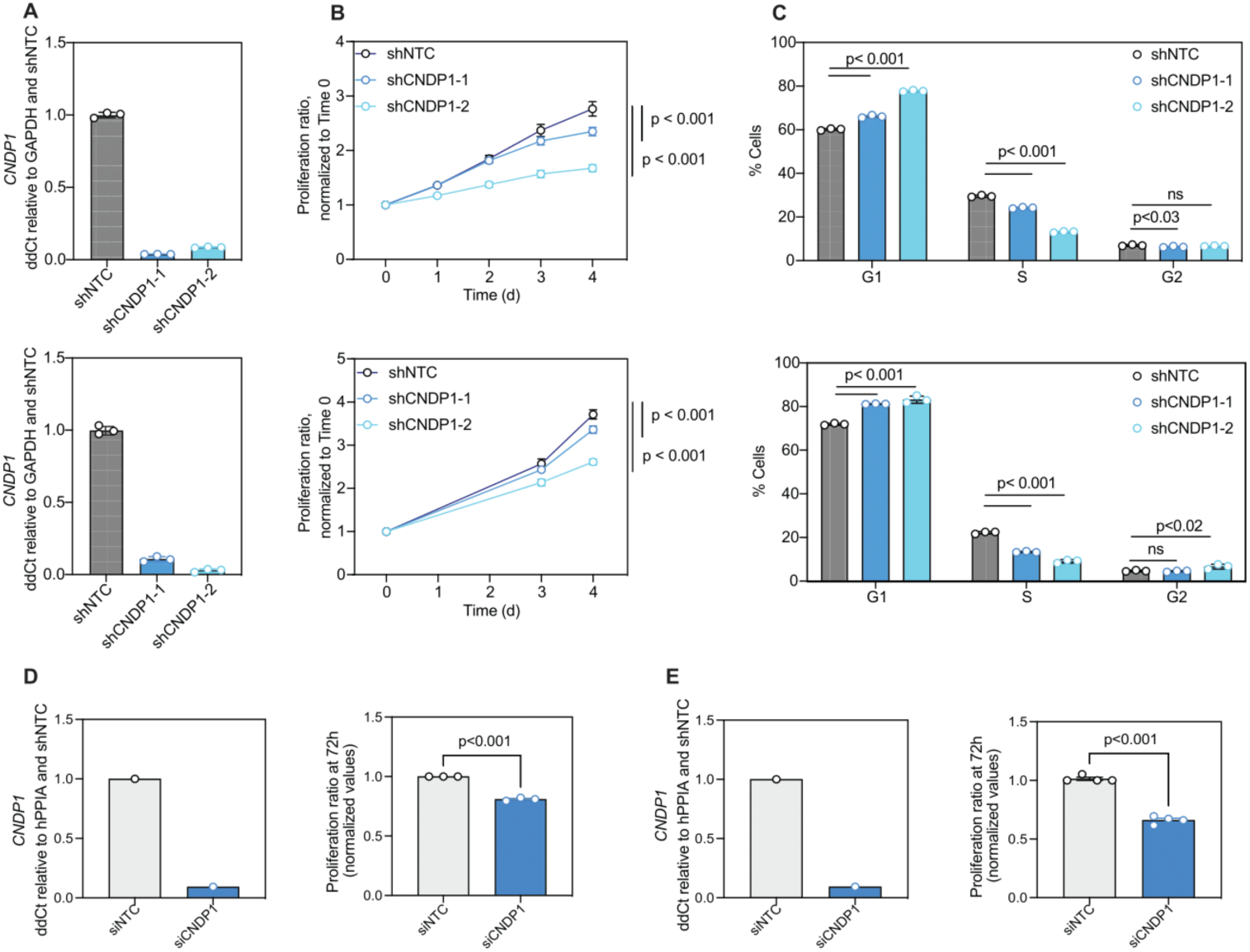
CNDP1 modulation impacts 2D proliferation of melanoma cells. **(A)** qRT-PCR quantification of *CNDP1* mRNA, relative to GAPDH and normalized to shNTC, in 10-230 BM (top) and 12-273 (bottom) cells expressing indicated shRNA. **(B) Top.** Proliferation curve performed by analyzing % confluency extracted from Incucyte image analysis normalized to day 0 of 10-230 BM cells expressing indicated tet-On shRNA (n=3, 96 h). **Bottom.** Proliferation curve performed by serial fixing and crystal violet staining of 12-273BM cells expressing indicated tet-On shRNA. Representative experiment shown of n=3 biological replicates. Statistics derived from one-way ANOVA testing between groups on the final time point. **(C)** Bar plots representing % cells distributed along cell cycle phases assessed by Edu differential staining (n=2) in 10-230 and 12-273 BM cells expressing indicated tet-On shRNA. Statistical analysis by one-way ANOVA with Dunnett multiple hypothesis testing correction. **(D) Left.** qRT-PCR quantification of *CNDP1* mRNA, relative to hPPIA and then normalized to siNTC, in 10-230 BM. **Right.** Proliferation ratio of 10-230 BM cells transfected with indicated siRNAs after 72 h of culture normalized to 0h (n=3), indicated as % confluency extracted from Incucyte image analysis. Multiple biological replicates represented. Statistical analysis by one-way ANOVA. **(E). Left.** qRT-PCR quantification of *CNDP1* mRNA, relative to hPPIA and then normalized to sINTC, in 12-273 BM. **Right.** Proliferation ratio of 12-273 BM cells transfected with indicated siRNAs after 96 h culture normalized to 0h (n=4), indicated as % confluency extracted from Incucyte image analysis technology. Multiple biological replicates represented. Statistical analysis by one-way ANOVA. See Figure S2

### CNDP1 inhibition suppresses brain metastasis establishment and maintenance in xenografts, and Cndp1 ectopic expression confers brain tropism to syngeneic models

To test if CNDP1 is required for BM establishment and/or growth *in vivo*, NSG mice were injected intracardially with patient-derived 12-273 BM cells stably transduced with either CNDP1-targeting or non-targeting control shRNA (Figure 3A-H). *In vivo* experiments were carried out using two approaches: i) constitutive CNDP1 knockdown, in which dox-containing chow was administered to mice right before tumor cell injection (Figure 3A-D), which addresses the requirement of CNDP1 for the *establishment* of metastasis, and ii) a more therapeutically relevant approach in which dox was administered after metastases were seeded, which assesses if CNDP1 is required for metastasis *maintenance* and growth (Figure 3E-H). Mice were humanely euthanized when any group started to show symptoms of distress or more than 20% weight loss, and organs were collected and processed for histological analyses. To quantify metastasis burden, we used both whole body bioluminescence live imaging (BLI) and MetFinder(50), a machine learning-powered tool we developed to segment, measure and quantify metastases in H&E-stained sections of murine tissues at termination. Our data shows strongly reduced brain and liver metastatic burden upon CNDP1 silencing with both shRNA (p<0.03, n.s. for shCNDP1-1 in the inducible approach in liver; Fig. 3D,H) A similar phenotype was observed in another patient-derived model, 10-230 BM, and in a brain-tropic breast cancer model (MDA-231 Brm2), after either constitutive (Fig S3A-D) or inducible CNDP1 knock-down (Fig S3E-L), supporting a role for this dipeptidase across cancer types. RNA-seq of GFP positive cells sorted from metastatic lesions confirmed that *CNDP1* silencing was maintained *in vivo* (Fig. S3M).

**Figure 3.**
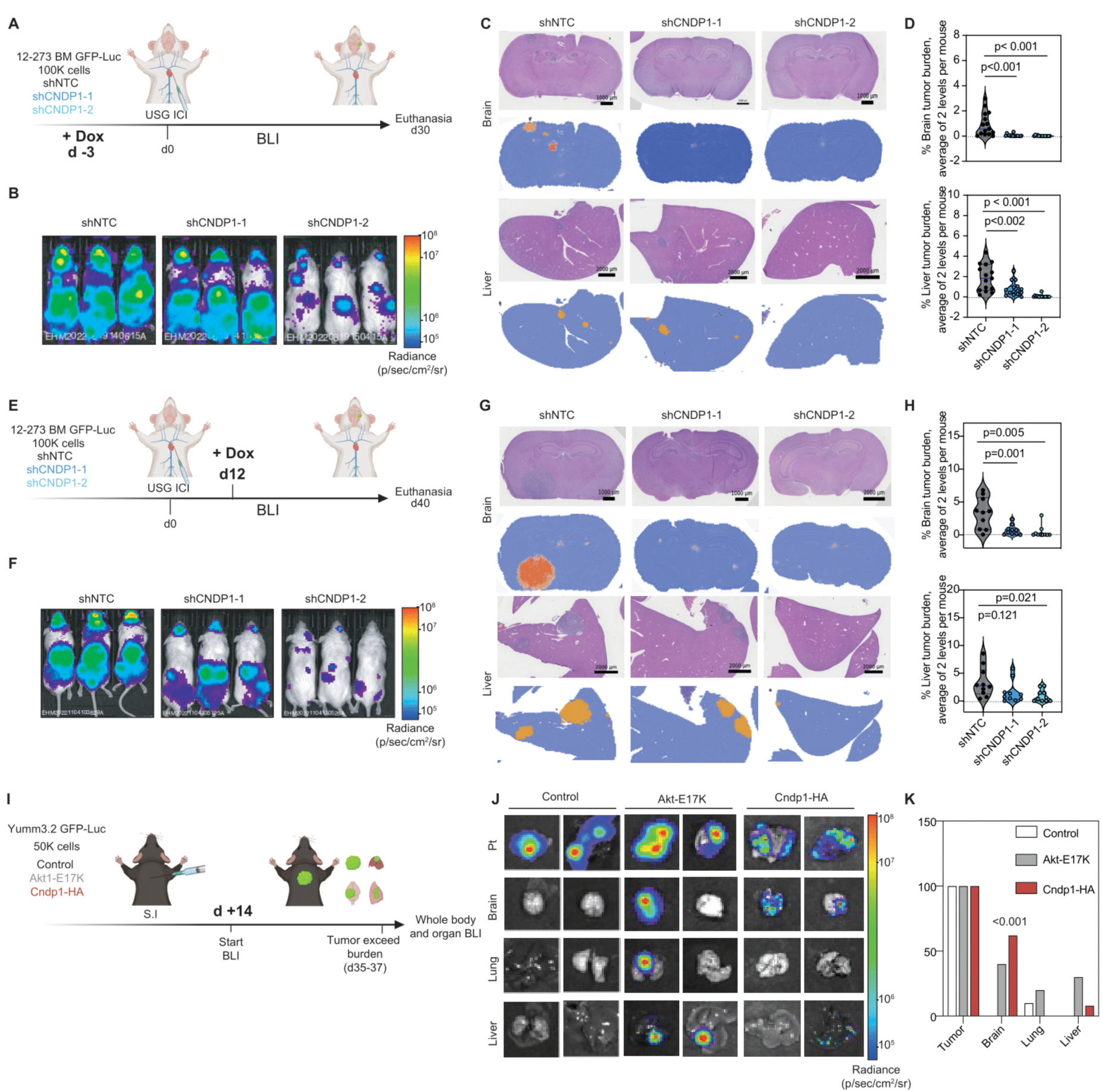
CNDP1 silencing impairs establishment and maintenance of *in Vivo* melanoma metastasis, and Cndp1 ectopic expression drives melanoma brain metastasis. **(A)** Schematic representation of the *in vivo* xenograft constitutive approach experiment (USG ICI: Ultra Sound Guided Intra-Cardiac Injection). **(B)** Representative images of whole-body *in vivo* BLI of NSG mice injected with 12-273 BM cells transduced with indicated tetON-shRNA 28 days post-injection (n = 12 mice per group, BLI represented by radiance: p/sec/cm^2^/sr). **(C)** Representative H&E images and corresponding MetFinder-generated heatmaps of brain and liver sections of NSG mice injected with indicated cells (n = 12 mice per group, two sections at 3 levels analyzed per organ/mice. Scale bars represent 1000 μm and 2000 μm for brain and liver, respectively). **(D)** Tumor burden in brain parenchyma, leptomeninges and liver quantified as the percent of area occupied by tumor and average of 2 levels of sectioning per organ (100 μM separation), obtained by MetFinder analysis. Statistical analysis by one-way ANOVA. **(E)** Schematic representation of the *in vivo* xenograft therapeutic approach experiment (USG ICI). **(F)** Representative images of whole-body *in vivo* BLI of NSG mice injected with 12-273 BM cells transduced with indicated tetON-shRNA at day 36 post-injection (n = 12 mice per group, BLI expressed as radiance: p/sec/cm^2^/sr). **(G)** Representative H&E and MetFinder brain and liver metastasis sections of NSG mice injected with indicated vectors (n = 12 mice per group; duplicated sections and levels analyzed per organ/mice. Scale bar representative of 1000 μm and 2000 μm for brain and liver, respectively) **(H)** Tumor burden in brain parenchyma, leptomeninges and liver quantified as the percent of tumor-occupied area vs total organ area and averaged of 2 levels of sectioning per organ. Obtained by performing MetFinder analysis. Statistical analysis by one-way ANOVA. **(I)** Schematic representation of *in vivo* allograft experiment with indicated conditions. S.I: Subcutaneous injection. **(J)** Ex vivo BLI of organs in the corresponding groups of mice injected with YUMM3.2 cells control or transduced with Akt1-E17K or Cndp1-HA in indicated organs (Pt=Primary tumor). (n = 9,10 &11 mice per group, respectively. BLI expressed as radiance: p/sec/cm^2^/sr). **(K)** Bar graph representing % of tumor incidence for primary (Pt), brain, lung and liver metastasis in mice injected with indicated conditions. Statistical analysis results from one-way ANOVA comparing metastasis incidence at end point (1=yes, 0=no). See Figure S3

Next, we tested whether CNDP1 ectopic expression is sufficient to enhance BM development. Murine Yumm3.2 melanoma cells (Braf^V600E/wt^ Cdkn2a-/-Pten-/-(51–53)), which do not metastasize when injected subcutaneously in C57BL/6J mice, were stably transduced with lentiviral vectors carrying HA-tagged Cndp1 or Akt1-E17K, which was previously shown to enhance metastasis development in this model(52) (Figure 3I). Cndp1 expression is not detected in these murine melanoma cells (not shown). Ectopically expressed Cndp1 exhibits carnosinase activity and confers a proliferation advantage *in vitro* (p<0.01, Figure S2E-G). Control, Akt1-E17K- and Cndp1-HA-transduced Yumm3.2 cells were subcutaneously injected in C57BL/6 mice. Upon reaching the maximum tumor volume, mice were euthanized, and organs harvested and examined by ex-vivo BLI for metastasis burden (Figure 3J-K). We observed that Cndp1-HA expressing cells gained exclusive brain tropism, with a higher incidence than Akt1-E17K transduced cells (62% vs 0 % (parental) vs 40% (Akt-E17K), p<0.001), which were also able to form lung and liver metastasis (Figure 3J-K). All these data points to a specific role for CNDP1 in enhancing BM development.

Our results support a role for CNDP1 in brain metastasis and expose CNDP1 targeting as a promising therapeutic intervention against these tumors. Interestingly, the observation that single nucleotide polymorphisms (SNPs) impairing CNDP1 secretion protect diabetic patients from developing nephropathy (36,54) has sparked interest in developing CNDP1 inhibitors, some of which have shown promising activity in preclinical models(55). Our studies provide a rationale for further developing small molecule compounds or antibodies targeting CNDP1.

### CNDP1 knockdown elicits an Integrated Stress Response and translational reprogramming

To further characterize the role of CNDP1 to tumor progression, we employed an unbiased approach integrating LC/MS metabolomics and bulk RNA-seq of shCNDP1-transduced cells induced with doxycycline for 72h (Figure 4A, Tables S5,6). We analyzed the resulting data with the MetaboAnalyst algorithm(56) (Table S7), which uses hypergeometric tests to identify enrichment of both transcript-based and mass spectra-based data in metabolic pathways. We found that “Aminoacyl tRNA-synthesis” was the most deregulated metabolic pathway in tumor cells upon CNDP1 suppression (Figure 4B, Table S7). To examine if these changes in tRNA synthesis may have implications for translation, we performed additional MS/MS (proteomics) analysis of 10-230 BM cells transduced with indicated shRNA (Table S8) and integrated these data with LC/MS (metabolomics) data (Table S5). This analysis revealed an accumulation of amino acids such as phenyl-alanine, arginine, tyrosine and leucine together with their respective aminoacyl tRNA-synthetases (Figure 4C) upon CNDP1 silencing.

**Figure 4.**
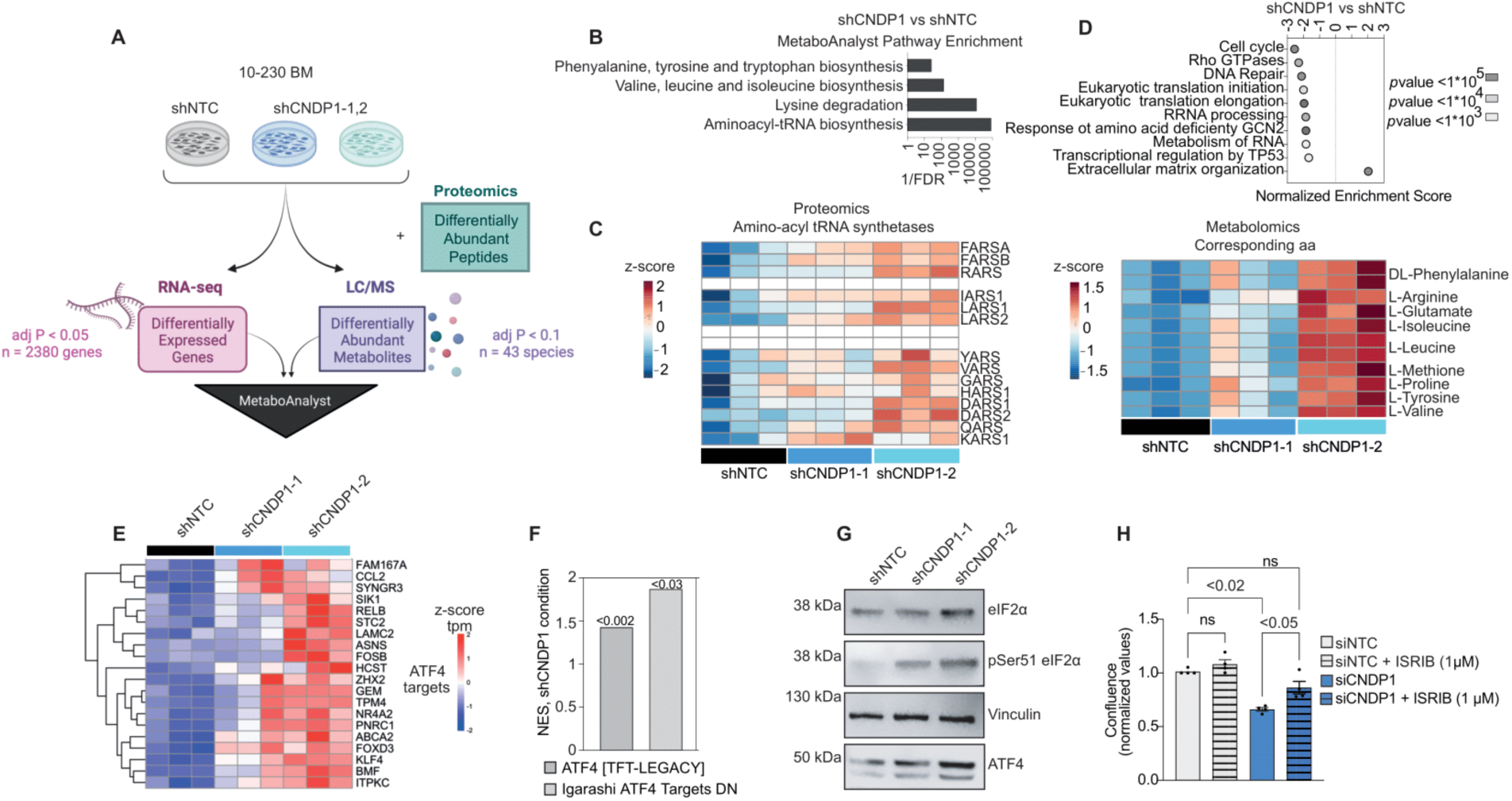
CNDP1 knockdown elicits an integrative stress response in melanoma cells. **(A)** Workflow of RNA-sequencing and LC/MS-based nonpolar metabolomics integrative studio in 10-230BM cells transduced with indicated tetON-shRNA. **(B)** Differential gene expression/metabolite abundance between shNTC vs shCNDP1-1 and shCNDP1-2 by indicated adjusted p values. MetaboAnalyst executed hypergeometric testing by combining p-values for metabolites and transcripts according to their relative proportion in each pathway. (RNA-seq: DESeq2, Metabolomics: T-test, pathways enrichment FDR < 0.05). See Tables S5-7 **(C)** Heatmap of t-Aminoacyl synthetases expression across different conditions paired with a heatmap of expression of their corresponding amino acids. MS/MS-based proteomics and LC/MS-based metabolomics were conducted on lysates of 10-230 BM cells expressing the indicated shRNA treated with doxycycline for 72 hours in 10% dialyzed serum DMEM. Differentially expressed proteins were identified by unpaired, two-tailed t-test comparing 2 groups normalized abundance: shCNDP1-1 and shCNDP1-2 vs shNTC (Data represented by z-score as indicated). **(D)** Integrated Gene Set Enrichment analysis (GSEA) of 12-273 and 10-230 BM shNTC vs both shCNDP1 tetON-shRNA RNA sequencing represented by Normalized Enrichment Score using Reactome Gene Ontology Analysis (RNA-seq: DESeq2, adjust p value < 0.001 and FC > 1.5). See Table S6. **(E)** Differential expression testing by DESeq2 nominated ATF4 targets in both shCNDP1 vs shNTC conditions in 10-230 BM RNA sequencing. (Data represented by z-score as indicated). **(F)** Pathway analysis via the Seq-n-Slide pipeline tested for pathway enrichment in differentially expressed genes between shCNDP1 and shNTC 10-230 BM RNA sequencing (Adjust p value indicated). **(G)** Representative immunoblots of phosphoSerine51 eIF2a, total-eIF2a, ATF4 and housekeeping (HK) protein Vinculin in lysates of 10-230 BM transduced with indicated tetON-shRNA. See Document S1 for details. **(H)** Proliferation ratio of 12-273 BM cells transfected with indicated siRNAs for CNDP1 silencing after 96 h of culture normalized to 0h (n=4), either treated with vehicle (DMSO) or ISRIB (phospho-eIF2a inhibitor, 1 uM) performed by analyzing % confluency extracted from Incucyte image analysis. Statistical analysis by one-way ANOVA. Proliferation ratio for 12-230 BM siNTC/siCNDP1 also shown in Figure 2 E. See also Figure S4

Further elucidating the cellular adaptative response triggered by CNDP1 silencing, transcriptomic and proteomics profiling revealed changes in translation initiation, elongation and RNA metabolism along with stress response pathways (Figures 4D, S4A, Tables S6,S9-10). Together, these changes were reminiscent of activation of the Integrated Stress Response (ISR), a cellular mechanism that halts or rewires protein synthesis in response to various sources of stress.(57) Indeed, we observed an increase in expression of ATF4 targets (Figure 4E,F) and p-Ser51-eIF2α levels (Figures 4G, S4B), both hallmarks of ISR activation, upon CNDP1 depletion. Moreover, a p-eIF2α inhibitor, ISRIB,(58) was able to partially rescue the proliferation defect induced by CNDP1 knock-down (p<0.02) (Figure 4H).

Although activation of the ISR often leads to a general shutdown of translation, CNDP1 silencing only had a modest impact on overall translation, in comparison to treatment with cycloheximide (CHX), a *bona fide* translation inhibitor(59) (Figure 5A). CNDP1 KD cells showed decreased mTOR activity, as revealed by p70 pS6K1, p-4E-BP1 and p-eIF4E levels, key modulators of translation initiation(60,61) (Figure 5B). mTOR modulation has been previously observed in response to TCA perturbation, such as a decrease in α-Ketoglutarate or changes in branched chain amino acid (BCAA) metabolism including isoleucine, leucine and valine, as observed upon CNDP1 KD (Figure 4B,C, Tables S5(62,63)). Additionally, transcriptomic data analysis of human melanoma metastases (i.e., TCGA, including various metastatic sites; GSE185386, including only MBM), revealed that *CNDP1* expression anti-correlates with translation and stress response gene signatures (Figure 5C). Our data pointed to a selective reprogramming of translation initiation rather than a complete shut down upon CNDP1 KD. To better understand specific translation changes elicited by CNDP1 KD, we performed polysome isolation followed by RNA-seq. Polysome profiling revealed changes in translation of mRNAs directly involved in mitochondria, ribosomes and cell division genes (Figure 5D-E, Figure S5A-B, Table S11). Concomitantly, we observed changes in eIF3d expression, suggestive of a shift from canonical to non-canonical cap-dependent translation possibly mediated by eIF3d/DAP5 (Figure S5C). It has been shown that eIF3d promotes selective cap-dependent translation of mRNAs involved in cell survival, such as mitochondrial genes (64,65). Of note, human BM cells expressing CNDP1^high^ levels are significantly enriched in mitochondrial genes in comparison to CNDP1^low^ cells (Table S12). Altogether, these data suggest that CNDP1 silencing initiates the ISR which reprograms the translation machinery to support mitochondrial respiration and cell cycle related transcripts.

**Figure 5.**
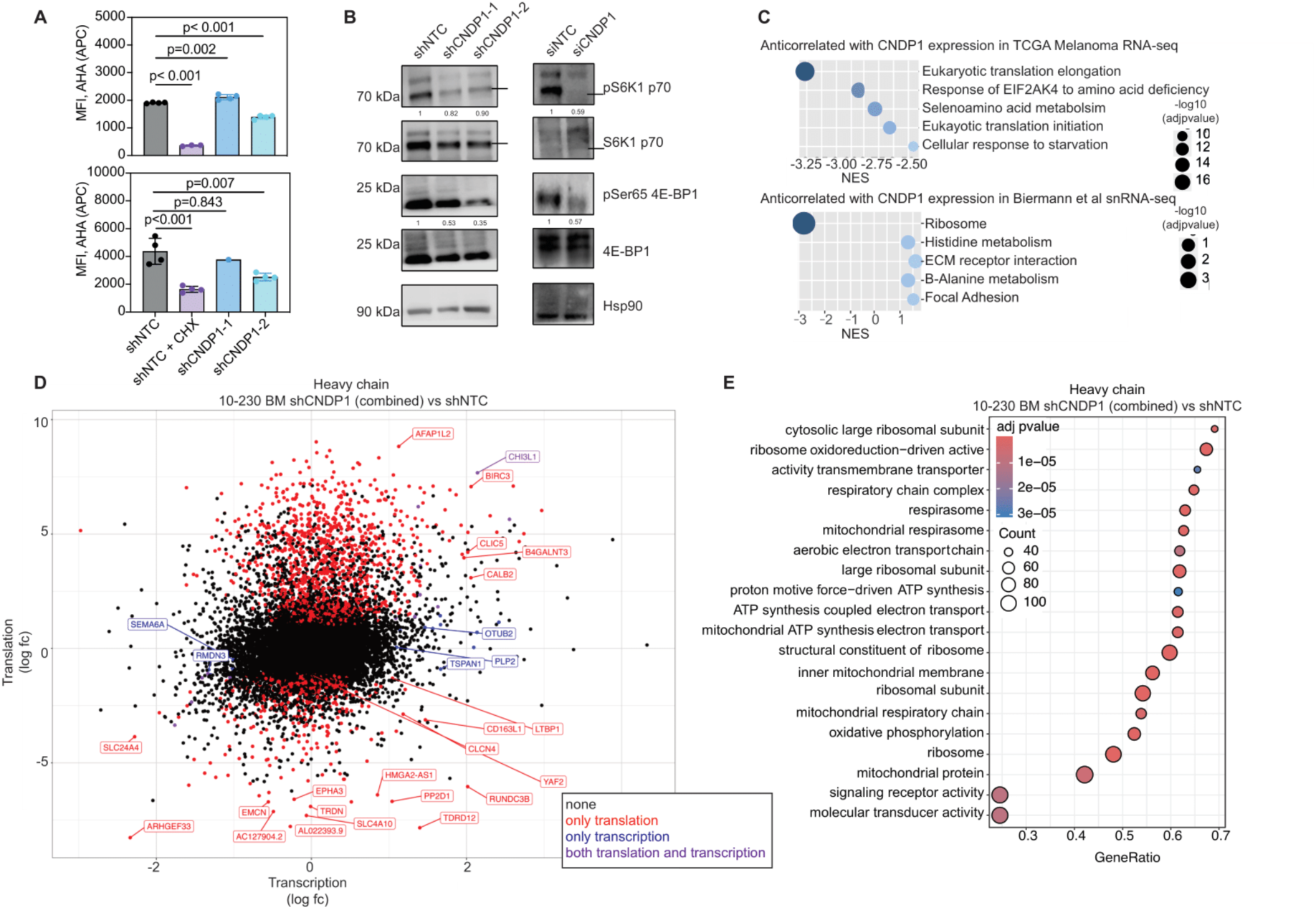
CNDP1 knockdown rewires translation program towards mitochondrial transcripts. **(A)** 10-230 and 12-273 BM transduced with indicated tetON-shRNAs and treated with doxycycline during 72h, labeled with AHA (APC) for 4 hours and analyzed by flow cytometry. Cycloheximide (CHX, 50 ug/ml) was used as a positive control of translation shutdown. Statistics rendered by one-way ANOVA with Dunnet’s multiple hypothesis testing correction. **(B) Left.** Representative immunoblots of phospho-S6K1 p70 (Threonine 389), S6K1 p70, phosphoSer65 4E-BP, total 4EBP and housekeeping (HK) protein Hsp90 in lysates of 10-230 BM transduced with indicated tetON-shRNA. **Right.** Representative immunoblots of phospho-S6K1 p70 (Threonine 389), S6K1 p70, phosphoSer65 4E-BP, 4EBP and housekeeping (HK) protein Hsp90 in lysates of 10-230 BM cells transfected with siRNA NTC or CNDP1 as indicated. Band correspondent to p70 S6K1 is indicated. Results for HRI, phosphoSer51 and total eiF2α expression from same samples shown in Figure 6K and Figure S4H, respectively. Ratio of protein phosphorylation shown normalized to total protein and HK protein levels. See Document S1 for details. **(C)** GENI (gene set enrichment identifier124) was applied to TCGA melanoma samples (n = 472) using the Reactome 2022 pathway database. Five anticorrelating gene sets with FDR < 0.05 were represented. **(D)** Genes differentially expressed in shCNDP1 (combined) vs shNTC polysome heavy chain (log fc Translation vs log fc Transcription; p<0.05, lfc > 1) at translation, transcription and both transcription and translation levels in 10-230 BM cells. See Table 11. **(E)** Integrated Gene Enrichment analysis of Heavy Chain RNA sequencing of 10-230 BM shCNDP1 (combined) vs shNTC represented by Normalized Enrichment Score using Reactome Gene Ontology Analysis (RNA-seq: adjust p value < 0.05). See also Figure S5

### CNDP1 depletion perturbs mitochondrial activity and results in increased intracellular copper levels

We next sought to understand how CNDP1 KD triggers mitochondrial stress, resulting in activation of the ISR and translation rewiring. Given our data indicating a shift towards translation of mitochondrial transcripts, we focused our attention on the mitochondria. Indeed, we observed profound morphological changes in the mitochondria of CNDP1 KD cells, including a smaller number of cristae per mitochondria (Figure 6A-E, Figure S6A-E), a phenotype previously reported as functionally defective(66)*. In vivo*, we observed reduced expression of the mitochondrial transport protein Tom20 in brain metastases upon acute CNDP1 KD (Figure 6F), which is indicative of dysfunctional mitochondrial structures (67). Taken together, these data raised the possibility that carnosine accumulation from CNDP1 inhibition causes mitochondrial dysfunction. Consistent with this hypothesis, both CNDP1 KD and carnosine treatment elicited a significant decrease in oxygen consumption rate (OCR) measured by Seahorse assays (Figure 6G, Figure S6F, p<0.02).

**Figure 6.**
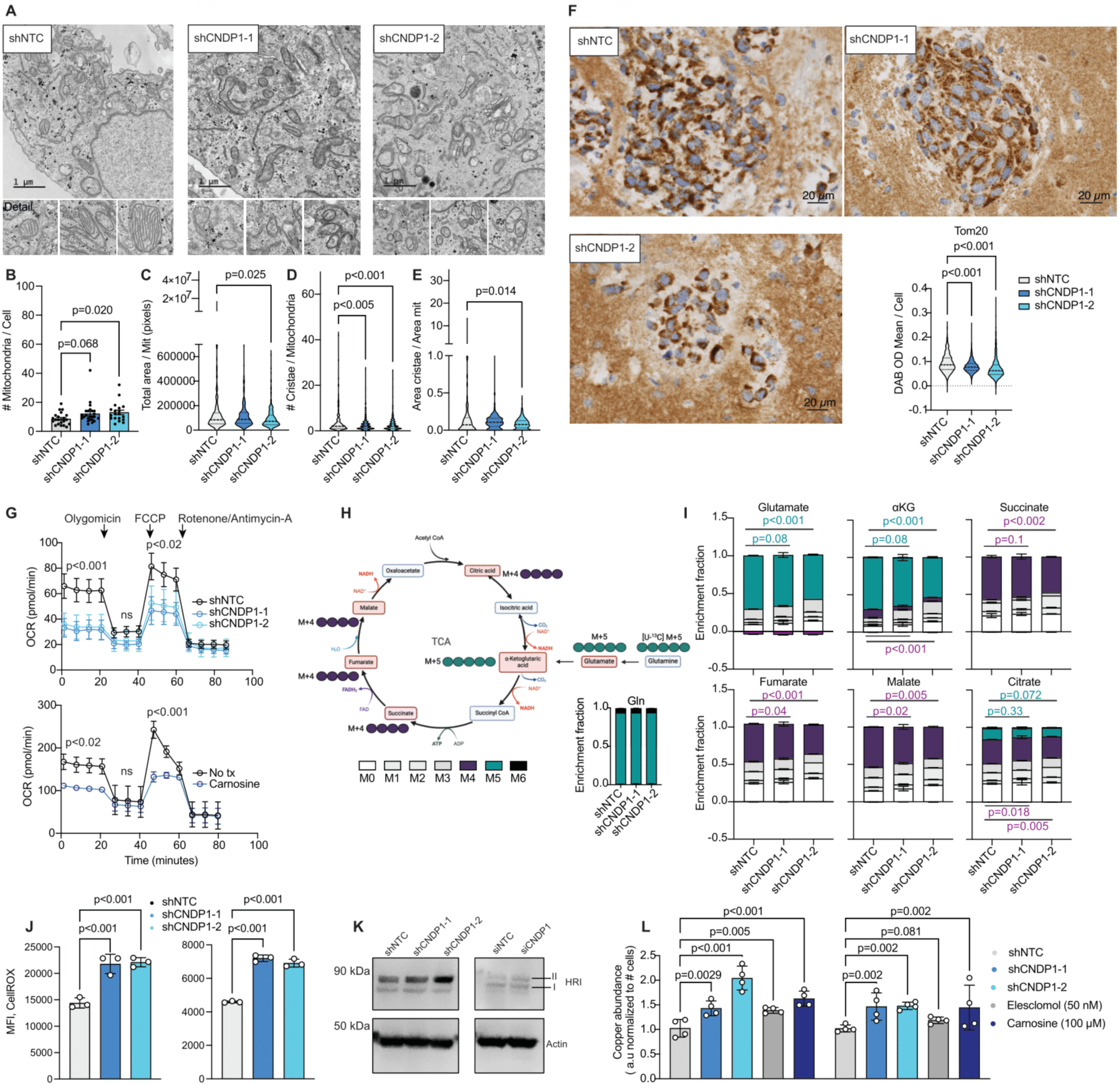
CNDP1 knockdown melanoma cells display profound mitochondrial defects, copper overload and ROS accumulation. **(A)** Mitochondria imaging using electron microscopy of 10-230 BM transduced with indicated tetON-shRNA. Scale =1 μM. **(B-E)** Quantification of number of mitochondria per cell **(B)**, mitochondria area **(C)**, cristae number per mitochondria **(D)** and cristae area per mitochondria **(E)** area in 10-230 BM. Statistical analysis by one-way ANOVA (p<0.05 or indicated). **(F)** Tom20 expression by IHC in BM from 12-273 BM cells transduced with indicated shRNAs induced with doxycycline during 5 days. Scale = 200 μM. Quantification of Tom20 cytosolic staining DAB intensity per cell and per mice. Statistical analysis by two-way ANOVA (p<0.001). **(G) Top.** Seahorse MitoStress analysis of OCR in 12-273 BM transduced with indicated tetON-shRNA. Statistical analysis by one-way ANOVA. Representative replicate shown (n=3, p<0.05 or indicated). **Bottom** Seahorse MitoStress analysis of OCR in 12-273 BM treated with Carnosine 40 mM overnight. Statistical analysis by one-way ANOVA. Representative replicate shown (n=3, p<0.05 or indicated). **(H)** Schematic model of glutamine metabolism in cancer cells. Green (M+5) and purple circles (M+4) represent carbons derived from [U-13C5] glutamine. The black arrows indicate oxidative and reductive carboxylation flux from glutamine. Cells were cultured with [U-^13^C5] glutamine for 6 h before metabolite extraction and gas chromatography-mass spectrometry (GC-MS) analyses. Normalized distribution of glutamine in mentioned conditions in 12-273 BM cells transduced with indicated tetON-shRNAs treated with doxycycline for 24h are shown. (n = 3); data are shown as the mean ± SD. **(I)** Normalized distributions of glutamine, glutamate, and TCA cycle metabolites, including α-ketoglutarate (αKG), succinate, fumarate, malate and citrate, in 12-273 BM cells transduced with indicated tetON-shRNAs treated with doxycycline for 24h are shown. (n = 3); data are shown as the mean ± SD. Statistical analysis by one-way ANOVA. P value indicated. **(J)** Mean fluorescence intensity of CellROX in live cells as determined by DAPI-status in 10-230 (left) and 12-273 BM (right) cells. Statistical analysis by one-way ANOVA. Representative replicate shown (n=3). **(K) Left.** Representative immunoblots of II & I HRI fractions and housekeeping (HK) protein b-actin in lysates of 12-273 BM transduced with indicated tetON-shRNA. **Right.** Representative immunoblots of II & I HRI fractions and housekeeping (HK) protein b-actin in lysates of 10-230 BM cells transfected with siRNA NTC or CNDP1 as indicated. II fraction considered as the autophosphorylated. Results for translation related proteins, phosphoSer51 and total eiF2α expression from same samples shown in Figure 5B and Figure S4H, respectively. Representative HK shown. See Document S1 for details. **(L)** Copper abundance represented as ratio of emission signal at 563 nm in indicated conditions in 10-230 BM (left) and 12-273 BM (right) normalized to total number of cells quantified by crystal violet (595 nm absorbance values). Statistical analysis by one-way ANOVA. Representative replicate shown (n=3). See also Figure S6

These findings led us to investigate if CNDP1 modulation also impacts Gln usage in the TCA cycle of melanoma cells. We performed Gln tracing experiments by using U-^13^ C5 labelled Gln and traced its incorporation by GC/MS analysis(68) in melanoma brain metastasis derived cells transduced with shRNA targeting CNDP1 as indicated (Figure 6H). We found that after 24h of silencing, CNDP1 KD cells display significantly reduced glutamine usage as evidenced by the lower levels of its glutamine-derived metabolites in the tricarboxylic acid (TCA) cycle: α-Ketoglutarate (M+5), Fumarate, Malate and Citrate (M+4) (Figure 6I). All these effects were mimicked by carnosine addition (Figure S6G-H). These data demonstrate that both CNDP1 KD and carnosine treatment impact TCA cycle activity. In addition to mitochondrial dysfunction, CNDP1 KD cells accumulated ROS, as indicated by CellROX measurements (Fig. 6J). Concomitant with ROS accumulation we observed the activation of heme-regulated inhibitor (HRI) kinase (Figure 6K), likely responsible for the above mentioned ISR activation (Figure 4).

Having observed that CNDP1 silencing impairs mitochondrial respiration and results in ROS accumulation, translation stalling and reprogramming, we looked for potential mechanisms directly triggering these effects downstream of CNDP1. Carnosine has been reported to work both as an ion chelator and ionophore, that is, able to transport iron and copper into the intracellular space.(69) We hypothesized that carnosine accumulation upon CNDP1 KD could lead to increased intracellular copper uptake through SLC transporters(70) and result in copper overload. Intriguingly, we observed CNDP1 KD cells exhibit dysregulation of genes involved in heavy metal metabolism such as iron (*LCN2, HMOX*) and copper transporters (*SLC46A3, ATP7B*) (Figure S6I-K). Indeed, by analyzing scRNA-seq data of MBM,(22) we found a significant enrichment in genes controlling copper homeostasis in CNDP1^high^ vs CNDP1^low^ MBM cells (Figure S6L). By staining cells with a permeable fluorescent dye (CS1) that binds free intracellular copper, we measured copper levels in CNDP1 KD and carnosine-treated cells along with cells treated with an established copper ionophore, Elesclomol(71), as a positive control. Indeed, both CNDP1 KD and carnosine-treated cells contain significantly higher intracellular copper levels, comparable to those of cells treated with Elesclomol (p<0.05 or indicated, Figure 6L). Copper mediated toxicity also affects the lipoylation of TCA enzymes such as Pyruvate dehydrogenase (PDH) and 2-oxoglutarate dehydrogenase (OGDH)(72), that results in degradation of these enzymes (Figure S6M), and Fe-S clusters assembly(73), leading to a decrease of both glutamine usage and OXPHOS dependent metabolism. Additionally, copper overload has been described to cause ROS accumulation(74). Therefore, intracellular copper overload in CNDP1 KD and carnosine-treated cells could explain the previously observed phenotypes of mitochondrial dysfunction and accumulation of ROS (Figure 6G-J, S6F,G).

Altogether, our studies support CNDP1 upregulation as a metabolic adaptation of cancer cells that increases their fitness during BM, likely by overcoming copper toxicity induced by carnosine accumulation in the brain metastatic niche (see model in **Figure S7**). CNDP1 inhibition emerges as a plausible strategy against metastatic progression by inducing metabolic stress.

## Discussion

Understanding distinct biological features of site-specific metastatic adaptation offers the possibility of discovering new targets and therapeutic options. In this study, we integrate two data sets, RNA-seq and proteomics,(25,26) comparing BM versus extracranial tumor samples to identify potential genes relevant to BM establishment and development. This analysis nominated a secreted dipeptidase, carnosine dipeptidase 1 or CNDP1, as a plausible player in melanoma brain metastasis. We also found *CNDP1* upregulated in lung and breast cancer brain metastases versus primary tumors across multiple databases. Silencing CNDP1 slows proliferation *in vitro* of brain metastasis-derived short-term cultures and impairs BM establishment and maintenance. A defect in liver metastasis upon CNDP1 silencing was also observed in our brain-tropic xenograft models; not surprisingly, since intracardiac injection favors hematogenous spread which allows our brain metastasis derived cells to yield not only brain but also liver metastases^25,116^. Moreover, carnosine is abundant in the liver^3^, thus CNDP1 upregulation may also provide a growth advantage there. However, Cndp1 ectopic expression exclusively conferred brain tropism to non-metastatic syngeneic models, supporting CNDP1 as a brain metastasis driver. In all, we demonstrate that high CNDP1 levels confer cancer cells increased ability to seed and colonize the brain. While CNDP1 target, carnosine, has been studied for its various chemical properties, the contribution of CNDP1 to human disease and in particular cancer is understudied. A mutant form of this dipeptidase was found associated with less severe nephropathy in type II diabetes patients(36,54); however, our study is the first to demonstrate the contribution of CNDP1 to brain metastasis.

Whether CNDP1 expression is induced once cells arrive to the brain, or if CNDP1^high^ cells have increased ability to seed in the brain parenchyma is unclear. In support of the latter, we found higher *CNDP1* levels in lymph nodes of melanoma patients that went on to develop brain metastases than in those that did not have BM. In melanoma, recent studies have shown cells can adopt different cell states, characterized by distinct transcriptional programs associated with varying metastatic potential(75,76). We did not find the expression of CNDP1 to be associated with specific transcriptional states (data not shown), yet we observed a plausible regulation of CNDP1 by MITF. Indeed, in Biermann et al., we reported an enrichment of the MITF signature in MBM vs extracranial metastasis.(22) However, our preliminary data suggest *CNDP1* upregulation in BM across non-melanoma tumors (i.e., breast and lung adenocarcinoma BM), likely by MITF-independent pathways. Further studies will be necessary to dissect CNDP1 regulation, and whether CNDP1^high^ cells can be detected in primary tumors and perhaps even hold prognostic value.

In this study, we propose that melanoma BM cells overcome carnosine-mediated copper overload in the metastatic niche by upregulating CNDP1. Copper uptake has been reported to enhance MAPK signaling in BRAFV600E mutant cells, and trigger MAPK inhibitor (MAPKi) resistance in melanoma cells(77). It also known how OXPHOS activation depends on MAPK signaling(78,79) . We therefore hypothesize that melanoma cells that resist copper overload may be able to not only escape from MAPKi treatment, but also to survive in the BM niche. By DepMap(80) data mining we found that MITF^high^ melanoma cell lines are more resistant to Elesclomol treatment (Figure S6N). Moreover, melanocytic-like (MITF^high^) cells may exhibit more tropism for BM(22) because of their well-known copper dependency, which these cells utilize for tyrosinase activity and melanin production(81–83) .

Our results reveal a detrimental effect of CNDP1 silencing in the establishment and maintenance of metastasis *in vivo* along with a mild effect on proliferation *in vitro* in our brain-derived models. The main reported substrate of CNDP1, carnosine, is known to be abundant in brain (e.g., 3 mg / 100 gr of tissue in rat brain), where it is produced mainly by oligodendrocytes and displays antioxidant, scavenging, pH buffering and anti-neuroinflammatory properties(38,41–43) . Carnosine has been studied as a dietary supplement in its native form, as well as in forms less easily cleaved by CNDP1, such as N-acetyl carnosine and zinc-L-Carnosine(34) . Small randomized controlled trials have shown that carnosine supplementation can benefit the short-term memory of healthy elderly patients, as well as those with Mild Cognitive Impairment(84,85). Our studies support previous evidence of carnosine’s ability to function as a copper ionophore(39) as part of a number of mechanisms that regulate tightly the distribution and transport of this micronutrient. These findings open a new avenue in the study of carnosine in copper-dependent mechanisms such as hepatic function or dopamine production and their potential effects in other diseases(81,82,86). For instance, highly abundant oligodendrocytes in the substantia nigra(87,88), that express both CARNS1 and CNDP1(49), may contribute to control carnosine-mediated copper uptake by dopaminergic neurons used for dopamine production. Meanwhile, carnosine supplementation has been reported to exert anti-proliferative effects on tumor cells(44). In highly glycolytic tumors, it has been reported that Isoform 2 of CARNS1 (CARNS2) controls carnosine levels, which have immunomodulatory effects during hepatic cancer progression(89). We hypothesized that carnosine accumulation could be detrimental to tumor growth, and that metastatic cells may benefit from cleaving carnosine and mitigating its antitumoral effects. The regulation of carnosine levels in tissues is an open question; for instance, humans are known to have higher levels of circulating CNDP1 in serum than other mammals(31,33). In fact, higher apes and the Syrian hamster have evolved an export signal sequence at the *CNDP1* locus, absent from other species, for unclear reasons(32,34). Carnostatine is a structural analog of carnosine that has been described to inhibit CNDP1 dipeptidase activity in preclinical models of diabetic nephropathy^52^. Future *in vivo* studies could test the therapeutic potential of pharmacological inhibitors of CNDP1 able to cross the blood brain barrier, alone or in combination with approved therapies.

CNDP1 is also known to cleave homocarnosine, a dipeptide made of GABA and L-histidine that exists at higher concentrations in the human than in the mouse brain(37,90,91). GABA is an inhibitory neuromodulator mainly produced in the cytosol of GABAergic neurons that regulates the activity of astrocytes and oligodendrocytes(92–94). We and others have reported that brain metastatic tumors can modulate the activity of neural cells in the metastatic niche by various mechanisms(25,95). In addition, GABA release from melanocytes has been shown to promote melanoma initiation by influencing their crosstalk with keratinocytes(96). Future studies could determine if increased GABA —from CNDP1-mediated homocarnosine cleavage — allows metastatic cells to hijack the neural microenvironment to their advantage.

To our knowledge, we are the first to report that CNDP1 silencing elicits translation reprogramming by modulating two interconnected pathways, ISR and mTOR, known to respond to various sources of cellular stress(61,97–100). ISR activation and mTOR inhibition upon CNDP1 KD result in preferred translation of mitochondrial and survival proteins, most likely controlled by eIF3d cap-dependent translation mechanisms(64,65), as previously shown under other forms of cellular stress. Extensive efforts have tried to determine if ISR activation is a barrier or conduit for tumor progression(29,99,101,102). ISR activation in response to CNDP1 KD leads to changes in the total AA pool, accumulation of tRNAs, and preferential translation of survival factors without inducing cell death. Genetic inhibition of CNDP1 significantly reduces brain metastasis burden; however, ISR activation upon CNDP1 KD could eventually allow cell survival in the metastatic niche. These adapted cells might be able to persist, acquiring additional changes that may permit tumor recurrence.

The shift of the translation machinery towards mitochondria-related transcripts paralleling copper accumulation upon CNDP1 KD particularly captured our attention. Whereas iron homeostasis is tightly post-transcriptionally regulated by Iron Response Elements (IREs)(103) and Iron Regulatory Proteins (IRPs)(103,104), similar mechanisms regulating copper metabolism have not yet been identified in human models(105–107). As discussed above, mitochondrial metabolism has been shown to play a key role in the establishment of brain metastasis (22,25,26). Proper oxidative phosphorylation relies on various factors,(21,22,26) including metal transport, a key limiting process that links mitochondrial function and ROS homeostasis(108). Recent studies have shown that cancer cells develop different adaptative mechanisms aimed at collecting heavy metals from the metastatic niche to support their metabolic function(109,110). Their reliance on heavy metals and particularly copper uptake becomes a vulnerability for brain metastatic cells. As such, upon CNDP1 silencing, accumulation of the ionophore carnosine (39,40) may induce copper toxicity(72,74,111) and mitochondrial malfunction. Supporting this model, CNDP1 depletion recapitulates the reported phenotypic effects of copper overload including ROS accumulation(40,74,112), reduced OCR(73) and degradation of lipoylated TCA enzymes(72,111).

Our work exposes the relevance of copper transport and uncovers a mechanism underlying oxidative phosphorylation reliance as a key metabolic adaptation to the brain metastatic niche. We found the dipeptidase CNDP1 tipping the balance of complex, interrelated mechanisms of cell survival emerging under metabolically demanding circumstances. Future studies will reveal how to further exploit these cell vulnerabilities to maximize the therapeutic potential of CNDP1 pharmacological inhibitors, alone or in combination with approved therapies.

## Methods

### Patient samples

Patients’ tumor samples from infiltrated lymph nodes were collected from melanoma patients who were prospectively enrolled in the Interdisciplinary Melanoma Cooperative Group Database (IMCG) at NYU Langone Health (IRB #10362). Inclusion criteria for this clinicopathological database included patients who presented to the Laura and Isaac Perlmutter Cancer Center (2002–2018) for melanoma and were accrued within two months of diagnosis. All patients signed NYU Langone Health Institutional Review board-approved consent to allow using their specimens for research and to conduct follow-up.

Tissue microarray samples were acquired by agreement with Dr. Levesque lab from University of Zurich. All patients included in this study have signed a patient release form, which has been approved by an ethics committee and assigned the numbers EK647 and EK800.

### Data mining

#### TCGA

Data from The Cancer Genome Atlas Skin Cutaneous Melanoma Firehose Legacy dataset (n = 472 samples, The Cancer Genome Atlas (RRID:SCR_003193) was analyzed using cBioPortal (RRID:SCR_014555), (113)), which reports RNA-sequencing data in RSEM. Survival analysis based on gene expression in the TCGA cohort was carried out using TIMER (RRID:SCR_018737, (114)), the Tumor Immune Estimation Resource, which draws Kaplan-Meier curves based on a user-defined cutoff of expression by percentage compared to the median, and reports p values based on log-rank testing. Primary and metastatic samples were analyzed separately.

#### Analysis of scRNA-seq data

Integration of samples from *Biermann et al.* (*22*) was carried out using Seurat’s CCA function (RRID:SCR_007322) in R studio (R Project for Statistical Computing (RRID:SCR_001905) and Bioconductor (RRID:SCR_006442). Tumor cells were parsed from non-tumor cells: non-tumor cells were excluded based on expression of genes unique to immune and stromal populations (using AddModuleScore and AUCell function (RRID:SCR_021327)). Next, malignant cells were segregated in *CNDP1* high and low using SCT (Single Cell Transform) (expression > 0 as threshold to plot a copper signature ((115)WP3286, WikiPathways (RRID:SCR_002134)) in *CNDP1*-high vs -low cells. CNDP1 expression from Non-Small Cell Lung Cancer primary and brain metastasis was obtained as described in *Tagore, Caprio et al.* in press in Nature Medicine.

#### Other databases

Raw, normalized or analyzed data were downloaded from Gene Expression Omnibus (GEO) (RRID:SCR_005012) following each study’s directions and code indicated in figure legends. Statistical analysis as described in figure legends.

#### TCGA-ATACseq

Normalized ATACseq counts from 370 TCGA patients were retrieved from UCSC Xena Browser database (RRID:SCR_018938). 10kb BIN segments around the *CNDP1* and *CNDP2* loci were visualized by heatmap using Prism10 (RRID:SCR_002798) and averaged for the ATACseq counts. Next, ATACseq counts in 10kb BIN segments around *CNDP1* promoters were averaged and plot against matched RSEM expression counts for Pearson’s correlation using Prism10 (RRID:SCR_002798).

#### GTEx, Genotype-Tissue Expression

We utilized Genotype-Tissue Expression (RRID:SCR_013042) data to assess expression from various tissues to identify tissue-specific and cross-tissue expression patterns of *CNDP1, CNDP2* and *CARNS*1. The data used for the analyses described in this manuscript were obtained from the GTEx Portal on 08/11/2023.

#### Depmap

Gene expression data for *MITF* was obtained from the “DepMap Public 22Q2” (Cancer Dependency Map Portal (RRID:SCR_017655)(80)) release, which includes RNA-seq data (TPM, transcripts per million) for melanoma cell lines. *MITF* expression data was filtered to retain melanoma-derived cell lines only by cross-referencing with metadata on cell line origin. For each melanoma cell line, the area under the dose-response curve (AUC) was extracted as a measure of Elesclomol sensitivity, with lower AUC values indicating higher sensitivity to the drug. Correlation was calculated using Pearson’s correlation coefficient.

### Plasmid generation

CMV-Luciferase-EF1α-copGFP (GFP-Luc) Lentivector Plasmid was purchased from BD Biosciences (BLIV511PA-1). Tet-pLKO-puro was purchased from Addgene (RRID:Addgene_98398). shRNAs were cloned as previously described 115) into Tet-pLKO-puro using AgeI (NEB, Cat#R3552S) and EcoRI (Thermo, Cat#FD0275) restriction sites. pLKO tet-on scrambled non-targeting control (shNTC) was purchased from Addgene (RRID:Addgene_47541).

The FG12-HA-AKT1^E17K^ Luciferase-lRES-eGFP lentiviral vector has been described(52). Cndp1-HA cDNA was synthesized by Gene Universal in the pENTR2B plasmid. Gateway cloning was used to recombine the pENTR2B-Cndp1-HA entry clone and the pDEST FG12-Luciferase-lRES-eGFP lentiviral destination vector (synthesized in Gene Universal) to generate the lentiviral vector FG12-Cndp1-HA-Luciferase-lRES-eGFP.

### siRNA pools and transfection

NON-targeting control (NTC) plus Human siRNA pools, ON-TARGET plus *CNDP1* and *MITF* siRNAs were purchased from Dharmacon (QTE-3158440G, L-005824-00-0005, L-008674-00-0005). 150,000 to 200,000 melanoma cells were seeded in 6-well plates and transfected with 20μM siRNA pools using Lipofectamine 2000 (Invitrogen,Cat#11668019) following manufacturer’s recommendations.

### Lentivirus production and infection

HEK293T (RRID:CVCL_0063) cells at 80% confluency were co-transfected with 12 μg of lentiviral expression constructs, 8 μg of psPAX2 (RRID: Addgene_12260) and 4 μg pMD2.G (RRID: Addgene_ 12259) vectors using Lipofectamine 2000 (Invitrogen, Cat#11668019) following manufacturer’s recommendations for Tet-pLKO-puro system. Packaging plasmids psPAX2 and pCMV-VSV-G (RRID: Addgene_8454) were used in combination with FG12-HA-AKT1-Ei7K Luciferase-lRES-eGFP (52,117) to generate virus for stable cell line generation as described below. At 48h and 72h post transfection, supernatants were collected, filtered (0.45 μm) and stored at −80°C. Melanoma cells were infected with lentiviral supernatants supplemented with polybrene (EMD Millipore, Cat#TR-1003-G) at a final concentration of 4μg/mL. Twenty-four hours after infection, cells were selected by cell sorting by GFP fluorescence for GFP-Luc transduced cells or treatment with puromycin (1 ug/mL, Thermo Fisher Scientific, Cat #A1113803), blasticidin (10 ug/mL, Invivogen, Cat#ant-bl-1) or hygromycin (300 ug/mL, Thermo, Cat#10687010) accordingly.

### Cell culture

Melanoma short-term cultures (STCs) were derived as described and used at passage (25,51,118). For maintenance cultures, STC cells were grown in DMEM (Corning, Cat #10-013-CM) with 10% fetal bovine serum (regular: Corning, Cat#35-010-CV, Tet-free: Neuromics, Cat#FBS002-T), 1 mM Sodium Pyruvate (Cytiva, Cat#SH30239.01), 4 mM L-Glutamine (Sigma-Aldrich, Cat #59202C), 25 mM D-Glucose (Gibco, Cat#A2494001), 1% Nonessential Amino Acids (Sigma-Aldrich, Cat#M7145-100ML), 100 units/mL penicillin, and 100 ug/mL streptomycin (Gibco, Cat# 15-140-163). For functional assays, cells were seeded in DMEM (Corning, Cat #10-013-CM) with 10% dialyzed Tet-free fetal bovine serum (Neuromics, Cat#FBS002-T), 4 mM L-Glutamine (Sigma-Aldrich, Cat #59202C) and 25 mM D-Glucose (Gibco, Cat#A2494001). HEK293T cells (RRID:CVCL_0063), MDA-231-Brm2 (RRID:CVCL_VR36) and SKMEL-239 (RRID:CVCL_6122) melanoma cells(119) were grown in DMEM (Corning, Cat #10-013-CM)with 10% fetal bovine serum (FBS, Corning, Cat#35-010-CV), 1 mM Sodium Pyruvate (Cytiva, Cat#SH30239.01), 4 mM L-Glutamine (Sigma-Aldrich, Cat #59202C), 25 mM D-Glucose (Gibco, Cat#A2494001), 100 units/mL penicillin, and 100 ug/mL streptomycin(Gibco, Cat# 15-140-163). YUMM3.2 cells were grown in F12/DMEM media (Thermo Fisher) supplemented with 10% FBS, 1 µg/mL penicillin-streptomycin (Thermo Fisher), and 1 µg/mL NEAA (Thermo Fisher). Cell lines were maintained in a 5% CO2 incubator at 37°C and were routinely tested for Mycoplasma contamination by using ATCC Kit (30-1012K) following manufacturer instructions.

### In vitro proliferation assay

3,500 to 4,000 cells per well were seeded in 96 wells and monitored for 72 to 96 hours. Cells transduced with Tet-pLKO-puro shRNAs were treated with doxycycline 1 ug/mL (Goldbio, Cat #24390-14-5) for at least 48 hours before seeding or with indicated treatment.

#### Incucyte

After allowing cells to adhere overnight, Incucyte Software V2022.1, 2022.2, and 2023.1 (Sartorius IncuCyte SX5 Live Cell Analysis System (RRID:SCR_026298)), were used to capture brightfield whole-well images. Masks for delineating confluence were manually adjusted to ensure only live cells were captured, such as a minimum of 100 μM size per object. Images were taken and analyzed every 6-12 hrs. Percent confluency was normalized to time=0 reading to obtain a fold change value.

#### Crystal violet staining

After allowing cells to adhere overnight, a baseline plate was obtained by removal of media and fixation in 0.1% glutaraldehyde for 15 minutes, then stored with PBS(Corning, Cat #21-031-CM) in each well at 4C. Remaining plates were fixed every 24 hours thereafter and/or at the experimental endpoint. Wells were stained with 0.5% crystal violet in PBS for one hour and washed extensively with water. Crystal violet retained by cells was then eluted by incubation in 250 μL 15% acetic acid for 1 hour with shaking, and absorbance at 590 nm was measured. Absorbances were normalized to the absorbances from the baseline plate to obtain a fold change value.

### Cell cycle analysis

Cell cycle analysis was carried out using the Click iT EdU kit (Thermo Fischer, Cat #C10636) to approximate DNA replication, and FxCycle Far Red (Thermo Fischer, Cat #F10348) to measure total DNA content. Briefly, STC cells expressing control or CNDP1-targeting Tet-On shRNA’s were plated at 250,000 cells per well in 6-well plates with 1 ug/mL doxycycline. An additional dose of doxycycline was added at 48 h post plating. At 72h post-plating, cells were incubated with 10 μM EdU for 4 h. Then, cells were collected by trypsinization, washed in 3 mL 3% BSA in PBS, spun at 1200 rpm for 4 mins, fixed in 250μL Fixation buffer (Biolegend, Cat# 420801) for 20 mins in the dark at room temperature, washed in 3 mL 3% BSA, spun again. Permeabilization and Click-iT reaction were executed according to kit instructions, followed by FxCycle Far Red staining and analysis on BD LSR II Flow Cytometer.

### In vitro treatments

Crystalline >99% L-carnosine (Sigma-Aldrich, Cat #C9625-25G) was dissolved into Media or PBS (see Cell Culture) at a concentration of 200 mM by rocking in a light-protected 50 mL conical tube. Solutions were pH adjusted to match base Media by adding HCl dropwise (∼pH 7.45), then filtered using a 0.2 μm filter. Stocks were light-protected and maintained at -20C until use, when they were thawed at room temperature and diluted into media. Cells were treated in a 10 μM to 50 mM range. ISRIB (phospho-eIF2a inhibitor, Selleck Chemicals, Cat#S7400) was diluted in DMSO at 1mM stock solution and added *in vitro* at 1 μM final concentration for 72 hours. Elesclomol (Selleck Chemicals, Cat#S1052) was diluted in DMSO 10 mM and added *in vitro* in a range of 10 nM to 50 μM for 24 hours.

### L-Histidine assay

Carnosinase activity was analyzed by measuring liberated L-histidine using a protocol adapted from Teufel et al 2003(33). Briefly, conditioned media was collected, centrifuged to remove dead cells and debris, and supernatant isolated. Adherent cells were rinsed in PBS and scraped, and pellets were lysed in 50 mM Tris pH 8 with protease inhibitor (Roche, Cat#11836145001) using a Bioruptor sonicator for 5 rounds of 1 min with 1 min off in between each cycle. Lysates were centrifuged at 20,000 rpm for 15 mins at 4C and supernatants collected. Lysates or conditioned media were diluted 1:1 (150 uL: 150 uL) in 50 mM Tris. 50μL of 20mM MnCl2 was added to each tube, and samples were incubated for 30 mins at 37C with gentle shaking. L-Histidine diluted in culture media was used as a positive control. 100μL of 50mM L-Carnosine in Tris was then added to each sample, followed by incubation at 37C for 120 min with gentle shaking. Finally, 500 μL of 10% trichloroacetic acid in 50 mM Tris was added to each sample. Samples were centrifuged at 3000 rpm for 10 mins to remove precipitated proteins. 100μL of each supernatant was transferred to fresh 5 mL conical tubes. In a dark chemical hood, 2mL 0.3N NaOH was added to each sample, followed by 400 μL of freshly made 5 mg/mL o-phthaldialdehyde solution (OPA) in 2 N NaOH. Solutions were allowed to incubate at room temperature for exactly 3 mins in the dark before addition of 400 μL 6M HCl, at which point samples were capped and shaken vigorously for 5s each. 100 μL of each sample was transferred into a well of a black bottom 96 well plate. L-Histidine diluted in culture media was used as a positive control. L-Histidine fluorescence was detected using excitation wavelength of 344 λ and emission wavelength of 420 λ.

### AHA integration translation assay

10-230 and 12-273 BM cells expressing control or CNDP1-targeting Tet-On shRNA’s. 72h after doxycycline treatment initiation, cells were labeled according to recommendations from the Click-iT AHA labeling kit (Invitrogen Cat, #C10102). Briefly, cells were pre-incubated in methionine-free, serum-free media (Fisher Scientific with repleted glutamine, Cat#21-012-026), with or without doxycycline/carnostatine/carnosine depending on group condition, with or without cycloheximide (Sigma-Aldrich, Cat #C7698-5G) for 30 mins. Then, cells were labeled with AHA at a 50 μM final concentration for 4 hrs. Cells were trypsinized and permeabilized in 100 μL saponin-based permeabilization buffer, washed in 1 mL 1% BSA in PBS, and incubated on 500 μL alkyne detection reaction cocktail with 5 μM Alexa Fluor 647 Alkyne for 30 mins at room temperature. After washing with 1 mL 1% BSA in PBS, cells were resuspended in 100 μL 1% BSA and stained with DAPI immediately prior to FACS analysis on a BD LSR II Flow Cytometer using the APC and DAPI channels to gate for singlets. AHA incorporation in place of methionine was quantified as the mean fluorescence intensity in the APC channel.

### Quantitation of Reactive Oxygen Species

STC cells expressing control or CNDP1-targeting Tet-On shRNA’s were plated at 250,000 cells per well in 6-well plates with 1 ug/mL doxycycline. An additional dose of doxycycline was added at 48h post plating. At 72h post-plating, cells were incubated by replacing media with 1 mL fresh serum-free media containing indicated doses of Tert-butyl hydrogen peroxide (TBHP, positive control, ROS-stimulating agent, Thermo Scientific, Cat#180340050) or N-Acetyl Cysteine (NAC, antioxidant, Sigma-Aldrich, Cat #180340050) for 2.5 hrs. Fresh resuspensions of THBP and NAC were made on the day of each experiment in deionized sterile water. Then, CellRox DeepRed reagent was added to each well (excluding negative controls) at a final concentration of 500nM and allowed to incubate with cells for 30 mins at 37C protected from light. Media was then collected, then adherent cells were rinsed with PBS, trypsinized, and added to their respective conditioned media. After centrifugation at 1200 rpm for 4 minutes, cells were resuspended in 100 μL 3% BSA and kept on ice until analysis on the BS LSRII Flow Cytometer. DAPI was added immediately prior to analysis to detect dead cells. Cells were analyzed on the cytometer in order of treatment group, not by shRNA expression group. Only live singlet cells were compared, using the Mean Fluorescence Intensity of APC channel signal to approximate ROS burden.

### Copper detection

20K cells were seeded in black wall clear bottom 96 well plates treated for cell culture. All conditions were treated with doxycycline and/or indicated Elesclomol and carnosine doses. After 24 treatment, Coppersensor-1 (CS1, MedChem Express, Cat#HY-141511) was utilized at 5 µM following manufacturer indications.

### RT-qPCR

RNA was isolated from cells at 80% confluence in 6 well plates using the RNeasy Mini Kit (Qiagen, Cat #74104) with DNase I treatment. 1 ug of RNA was subjected to reverse transcription following TaqMan™ Reverse Transcription Reagents manufacturer indications (Applied Biosystems, Cat #4304134). Real time quantitative PCR was performed in 384 wells by mixing Applied Biosystems™ Power SYBR™ Green PCR Master Mix (Applied Biosystems, Cat #4367659), forward and reverse target gene primers at 0.2 nM final concentration (See Materials Table) and 100 to 150 ng of cDNA up to 15 μl final volume.

### Immunohistochemistry

Immunohistochemistry was performed using mouse anti-human CNDP1 Monoclonal (OTI1A5, LSBio) and Tom20 (D8T4N) Rabbit mAb (Cell Signaling Technology) on formalin fixed, paraffin embedded tissues. In brief, after deparaffinization and rehydration, heat-induced epitope retrieval was performed in 0.01 M citrate buffer (pH 6.0) in a 1200-W microwave oven at 100% power for 20 min. Sections were then cooled in tap water for 5 min, quenched in hydrogen peroxide for 30 min, washed with PBS, and incubated with blocking serum for 30 min followed by each primary antibody (Ab) diluted in buffer at room temperature for 1 h and at 4 °C overnight. Abs used were Tom20 (D8T4N) Rabbit mAb (Cell Signaling Technology, Cat #42406, RRID:AB_2687663) and CNDP1 Mouse anti-Human Monoclonal (OTI1A5) ab (LSBio, Cat#LS-C172361, RRID: AB_3676497). Slides were washed in buffer and incubated with diluted biotinylated for Tom20 (Sigma-Aldrich, Cat#A0545, RRID:AB_257896) or alkaline phosphatase for CNDP1 secondary antibodies for 1 h (Alkaline Phosphatase Ab mouse, LSBio, LS-C44264, RRID:AB_1050644). Avidin-biotinylated horseradish peroxidase complexes diluted at 1:500 were added, and complexes were visualized with diaminobenzidine for Tom20 staining or with Fast Red (Abcam, Cat# ab64254) for CNDP1 following manufacturer indications. Slides were then washed in distilled water, counterstained with hematoxylin, dehydrated, and mounted with permanent media. Appropriate positive and negative controls were included with the study sections.

### Western Blot Analysis

Cells were harvested in cold RIPA lysis buffer (Pierce, Cat # 89900) supplemented with protease inhibitor and protein was quantified by DC assay. Cell lysates (30 ug of protein) were resolved in 4%-12% Bis-Tris gels (Thermo Fisher Scientific, Cat#NP0321BOX) and transferred to PVDF membranes using wet transfer.

Membranes were blocked in either 5% non-fat milk or 5% BSA in Tris-buffered saline/1% Tween 20 (TBST) for 1 hour. Membranes were incubated overnight with primary antibodies (Ab) diluted in 5% milk or 5% BSA TBST. Primary ab used were anti-Human CNDP1 (352216) Ab (Novus Biological, Cat#MAB2489-SP, RRID: AB_3676498), PhosphoeIF2α (Ser51) Ab (Cell Signaling Technology Cat#9721S, RRID: AB_330951), EIF2S1 Monoclonal Ab (EIF2-alpha) (Invitrogen, Cat#AHO0802, RRID: AB_2536316), Monoclonal Anti-Vinculin Ab produced in mouse (Sigma-Aldrich, Cat#V4505-100UL. RRID: AB_477617), Atf 4 (D4B8) Rabbit mAb (Cell Signaling Technology, Cat#11815S, RRID: AB_2616025), Phospho-p70 S6 Kinase (Thr389) (108D2) Rabbit mAb (Cell Signaling Technology, Cat#9234T, RRID:AB_2269803), p70 S6 Kinase Ab (Cell Signaling Technology, Cat#9202, RRID: AB_AB_331676) 4E-BP1 (53H11) Rabbit mAb (Cell Signaling Technology, Cat#9644T, RRID:AB_2799054), Phospho-4E-BP1 (Ser65) Ab (Cell Signaling Technology, Cat#9451T, RRID: AB_330947), Hsp90 Ab (Cell Signaling Technology, Cat#4874S, RRID: AB_2121214), Rabbit HRI Polyclonal Ab (MyBioSource.com, Cat#MBS2538144, RRID:AB_2885168), Monoclonal Anti-beta-Actin-Peroxidase Ab produced in mouse, clone AC-15, purified from hybridoma cell culture (Sigma-Aldrich, Cat#A3854-200UL, RRID:AB_262011, Rabbit anti-eIF3D/EIF3S7 Ab, Affinity Purified (Bethyl Laboratories, Cat#A301-758A, RRID:AB_1210970, Eif4E Mouse, Unlabeled, Clone: 87, (BDCat#610270, RRID:AB_397665), eIF4G Ab (Cell Signaling Technology, Cat#2498, RRID: AB_2096025), DAP-5 Ab (F-2) (Santa Cruz Technology, Cat#sc-137011, RRID:AB_2095908 Pyruvate Dehydrogenase (PDH) Ab (Cell Signaling Technology, Cat#2784, RRID: AB_2162928), 2-oxoglutarate dehydrogenase (OGDH) Ab (Cell Signaling Technology, Cat#26865, RRID:AB_2737585), Anti-HA-Tag Rabbit Monoclonal Ab (Cell Signaling Technology, Cat #C29F4, RRID:AB_10693385). Membranes were washed 3 times for 10 min in TBST and incubated in secondary antibody for 1 hour in 5% milk or BSA TBST. Membranes were washed 3 times for 10 minutes in TBST and incubated with ECL substrate for 1 min and exposed to film or BioRad chemiluminescence dock station for imaging. Antibodies for phosphorylated proteins and their respective antibodies for total protein were blotted in different membranes or phosphorylated antibodies were blotted first, membrane was stripped and then the total was blotted. Representative housekeeping shown.

The following secondary antibodies were used: Goat anti-Rabbit IgG-Peroxidase (Sigma-Aldrich, Cat#A0545, RRID:AB_257896), Goat anti-mouse IgG kappa-light chain (Santa Cruz Technology, Cat#sc-516102, RRID:AB_2687626) 1:10,000.

Quantification was performed by inverting the membrane raw image, measuring the integrated density of each band using the same quantification area for all bands for the housekeeping bands and the protein of interest. Quantification is represented as a ratio of the signal of the protein of interest and the housekeeping protein signal and finally normalized to each control condition.

### Carnosine ELISA

Media (see Cell Culture above) was conditioned with melanoma cells for 24-72 hours (until reaching ∼80% confluency) and spun at 1200 rpm for 4 minutes to remove cells and debris. Carnosine ELISA was performed following manufacturer’s indications (Novus Biologicals, Cat#NBP2-75013).

### Metabolomics

#### LC-MS/MS with the hybrid metabolomics method

Samples were subjected to an LCMS analysis to detect and quantify known peaks. A metabolite extraction was carried out on each sample based on a previously described method(120). The LC column was a Millipore™ ZIC-pHILIC (2.1 x 150 mm, 5 μm) coupled to a Dionex Ultimate 3000™ system and the column oven temperature was set to 25 °C for the gradient elution. A flow rate of 100 μL/min was used with the following buffers; A) 10 mM ammonium carbonate in water, pH 9.0, and B) neat acetonitrile. The gradient profile was as follows; 80-20% B (0-30 min), 20-80% B (30-31 min), 80-80% B (31-42 min). Injection volume was set to 2 μL for all analyses (42 min total run time per injection).

MS analyses were carried out by coupling the LC system to a Thermo Q Exactive HF™ mass spectrometer operating in heated electrospray ionization mode (HESI). Method duration was 30 min with a polarity switching data-dependent Top 5 method for both positive and negative modes. Spray voltage for both positive and negative modes was 3.5 kV and capillary temperature was set to 320 °C with a sheath gas rate of 35, aux gas of 10, and max spray current of 100 μA. The full MS scan for both polarities utilized 120,000 resolution with an AGC target of 3e6 and a maximum IT of 100 ms, and the scan range was from 67-1000 m/z. Tandem MS spectra for both positive and negative mode used a resolution of 15,000, AGC target of 1e5, maximum IT of 50 ms, isolation window of 0.4 m/z, isolation offset of 0.1 m/z, fixed first mass of 50 m/z, and 3-way multiplexed normalized collision energies (nCE) of 10, 35, 80. The minimum AGC target was 1e4 with an intensity threshold of 2e^5^. All data were acquired in profile mode.

The resulting ThermoTM RAW files were converted to SQLite format using an in-house python script to enable downstream peak detection and quantification. The available MS/MS spectra were first searched against the NIST17 MS/MS (121), METLIN (RRID:SCR_010500) (122)and respective Decoy spectral library databases using an in-house data analysis python script adapted from our previously described approach for metabolite identification false discovery rate control (FDR) (123,124). Metabolite peaks were extracted based on the theoretical m/z of the expected ion type, e.g., [M+H]+, with a 15 part-per-million (ppm) tolerance and a ± 0.2 min peak apex retention time tolerance within an initial retention time search window of ±0.5 min. For all the group-wise comparisons, t-tests were performed using the Python SciPy SciPy (RRID:SCR_008058), (1.5.4) (125) library to test for differences and generate statistics for the downstream analyses. For the pairwise t-tests, any metabolite with a p-value < 0.01 was considered significantly regulated (up- or down-) for prioritization in the subsequent analyses. Heatmaps were generated by hierarchical clustering, performed based on the imputed matrix values utilizing R studio (R Project for Statistical Computing (RRID:SCR_001905) and Bioconductor (RRID:SCR_006442), (1.0.12). GraphPad Prism 9 (9.4.1, GraphPad Software GraphPad Prism (RRID:SCR_002798).

#### Data analysis

Metabolite peak intensities were extracted according to a library of m/z values and retention times developed with authentic standards. Intensities were extracted with an in-house script with a 10 ppm tolerance for the theoretical m/z of each metabolite, and a maximum 30 sec retention time window.

### RNA-sequencing analysis

After 72 h induction of shRNA expression with doxycycline, cells were scraped and RNA was harvested using the RNeasy Mini Kit (Qiagen, Cat #74104). For RNA-seq in cells from *in vivo* experiments, tissues were harvested after intracardiac injection and 7 days of doxycycline treatment. Tissues were homogenized by using a mechanic/enzymatic protocol (Collagenase Sigma, Cat#C2674-100MG and DNAse I, EMD Millipore, 260913-10MU) and sorted for GFP positive expression with adequate controls. RNA was extracted using the abovementioned kit. RNA-Seq library preps were made using the Illumina TruSeq Stranded total RNA with RiboZero Gold kit on a Beckman Biomek FX instrument, using 100 ng of total RNA as input, amplified by 12 cycles of PCR, and run on an Illumina NovaSeq 6000. Reads were aligned and gene expression compared using the Seq-n-Slide pipeline (Seq-N-Slide (RRID:SCR_021752)). Briefly, FastQC v0.11.7 was used to check fastq files for poor sequencing quality; all samples had high quality. Illumina adapter sequences and poor-quality bases were then trimmed using trimmomatic v0.36. Trimmed sequences were mapped to mm10 using STAR v2.6.0a (RRID:SCR_004463), indexed using samtools v1.9, then quantified for UCSC genes using htseq-count (RRID:SCR_011867) v0.11.1. Comparative analysis between conditions was performed using DESeq2 (RRID:SCR_015687) v1.24.0 with default parameters.

### Polysome Sequencing

Polysome isolation was performed by separation of ribosome-bound mRNAs via sucrose gradient as previously described(126). Briefly, 14X89mm Beckman Ultra-Clear centrifuge tubes were loaded with 5.5 ml of 50% and 5.5 ml 15% sucrose in Polysome Lysis Buffer (PLB) [20 mM Tris (pH 7.4) in Nuclease Free H2O, 10 mM NaCl, and 3 mM MgCl2 supplemented with RiboLock RNase inhibitor (ThermoFisher, Cat#EO0384) and 100μg/ml cycloheximide (Sigma-Aldrich, Cat #C7698-5G) and incubated at 4 °C horizontally overnight. 10-230 BM and 12-273 BM expressing shNTC and shCNDP1-1,2 were pre-treated with 100 μg/ml cycloheximide (Sigma-Aldrich, Cat #C7698-5G) for 15 min at 37oC, washed twice in phosphate-buffered saline (PBS) containing 100 μg/ml CHX, pelleted, and flash frozen (10% was separated for total RNA and protein analysis). Pellets were allowed to thaw on ice and resuspended in 750 μl of PLB. After 5 min of incubation on ice, 250 μl of detergent buffer (1.2% Triton, and 0.2 M sucrose in polysome isolation buffer) was added. Cells were further lysed for 10 min and lysate was clarified. Equal amount of clarified lysate was layered onto 15–50% sucrose gradients and sedimented at 36 000 rpm for 2 h in a SW41 rotor (Beckman Coulter) at 4 °C. Gradients were collected in 12 × 750 μl fractions by pumping 60% sucrose into the bottom of the gradient and collecting from the top using an ISCO fraction collector while simultaneously monitoring absorbance at 254 nm. RNA was isolated by extraction using TRIZOL (ThermoFisher, Cat# 15596026) as per manufacturer’s instructions. Light (poorly translated) polysome fractions (2–3 ribosomes) and well-translated heavy polysome fractions (≥4 ribosomes) were pooled. In parallel total RNA was extracted to normalize for transcriptional changes. All experiments were performed in triplicate. RNA quality and amount were determined by the Agilent Technologies kit and Nanodrop. RNAseq was carried out by the NYU School of Medicine Genome Technology Core using the Illumina Hi-Seq 2500 Single Read. Multiple triplicate samples were pooled for 10-230 BM. PCA analysis was performed using the “stats” R package (Bioconductor (RRID:SCR_006442)) and multiple individual triplicates (12-273 BM shCNDP11-1 and shNTC-2 bulk) were identified as outliers and removed from further processing.

Polysome data was analyzed using RIVET (RRID:SCR_026488, a R-Shiny-based tool(126). Utilizing the matched bulk and light + heavy polysome data, RIVET uses the “limma” R statistical package (RRID:SCR_010943) to determine transcripts that were over or under expressed in both transcription and translation. Within the RIVET app, the transcripts for both shRNAs targeting CNDP1 were combined to create a “combined” sample. The differentially expressed genes identified by RIVET were exported for further analysis. Using the R package “enrichplot,” GSEA (Gene Set Enrichment Analysis (RRID:SCR_003199)) analyses were performed on the differentially expressed gene lists to identify associated pathways possibly affected by the changed genes.

### Proteomics workflow

#### Protein isolation

10-230BM cells expressing shNTC, shCNDP1-1, or shCNDP1-2 were seeded at 1e6 cells per 15cm plate in triplicate and incubated in DMEM + 10% dialyzed serum + 1 ug/mL doxycycline for 72 hrs. STCs at ∼80% confluence were scraped from 10 cm plates on ice, washed once with cold PBS, and scraped in 1mL additional cold PBS into an eppendorf tube. After centrifugation at 1200 rpm for 4 minutes, pellets were flash-frozen in liquid nitrogen. Upon thawing, pellets were lysed in cold RIPA buffer supplemented with protease inhibitor, for 15 minutes with vortexing every 5 minutes. Protein concentration was determined using Micro BCA Protein Assay Kit (Thermo Scientific, Cat #23235).

#### Sample preparation for mass spectrometry analysis

150 μg of each protein lysate were proteolytically digested and subjected to quantitative mass spectrometry on an Orbitrap Fusion Lumos mass spectrometer using isobar tandem mass tags (TMT). Samples were prepared using the filter-aided sample preparation (FASP) method. Briefly, each sample was reduced with DTT (final concentration of 20 mM) at 57°C for 1 hour and loaded onto a MicroCon 30-kDa centrifugal filter unit (Millipore) pre-equilibrated with 200 μl of FASP buffer [8 M urea and 0.1 M tris-HCl (pH 7.8)]. Following three washes with FASP buffer, lysates were alkylated on a filter with 50 mM iodoacetamide for 45 min in the dark. Filter-bound lysates were washed three times each with FASP buffer followed by 100 mM ammonium bicarbonate (pH 7.8). The samples were digested overnight at room temperature with trypsin (Promega) at a 1:100 ratio of enzyme to protein. Peptides were eluted twice with 100 μl of 0.5 M NaCl. The tryptic peptides were subsequently desalted using an UltraMicro Spin Column, C18 (Harvard Apparatus) and concentrated in a SpeedVac concentrator.

#### TMT labelling

The dried peptide mixture was re-suspended in 100 μl of 100 mM Triethylammonium bicarbonate (TEAB) (pH 8.5). Each sample was labeled with TMT reagent according to the manufacturer’s protocol. In brief, each TMT reagent vial (0.8 mg) was dissolved in 41 μL of anhydrous ethanol and was added to each sample. The reaction was allowed to proceed for 60 min at room temperature and then quenched using 8 μL of 5% (w/v) hydroxylamine. The samples were combined at a 1:1 ratio and the pooled sample was subsequently desalted using strong-cation exchange and strong-anion exchange solid-phase extraction columns (Strata, Phenomenex) as described.

#### Global Proteome Analysis

A 500 μg aliquot of the pooled sample was fractionated using basic pH reverse-phase HPLC (as described) 250. Briefly, the sample was loaded onto a 4.6 mm × 250 mm Xbridge C18 column (Waters, 3.5 μm bead size) using an Agilent 1260 Infinity Bio-inert HPLC and separated over a 90 min linear gradient from 10 to 50% solvent B at a flow rate of 0.5 ml/min (Buffer A = 10 mM ammonium formate, pH 10.0; Buffer B = 90% ACN, 10 mM ammonium formate, pH 10.0). A total of 120 fractions were collected and non-concatenated fractions combined into 40 final fractions. The final fractions were concentrated in the SpeedVac and stored at -80°C until further analysis.

#### LC-MS/MS analysis

An aliquot of each final fraction was loaded onto a trap column (Acclaim® PepMap 100 pre-column, 75 μm × 2 cm, C18, 3, 100 Å, Thermo Scientific) connected to an analytical column (EASY-Spray column, 50 m × 75 μm ID, PepMap RSLC C18, 2 μm, 100 Å, Thermo Scientific) using Easy nLC 1000 (Thermo Scientific) with solvent A consisting of 2% in 0.5% acetic acid and solvent B consisting of 80% acetonitrile in 0.5% acetic acid. The peptide mixture was gradient eluted into the Orbitrap Lumos Fusion mass spectrometer (Thermo Scientific) using the following gradient: 5%-23% solvent B in 100 min, 23% - 34% solvent B in 20 min, 34% - 56% solvent B in 10 min, followed by 56%- 100% solvent B for 20 min. Full scans were acquired with a resolution of 60,000 (@ m/z 200), a target value of 4e5 and a maximum ion time of 50 ms. After each full scan the most intense ions above 5E4 were selected for fragmentation with HCD using the “Top Speed” algorithm. The MS/MS were acquired in the Orbitrap with a resolution of 60,000 (@ m/z 200), isolation window of 1.5 m/z, target value of 1e5, maximum ion time of 60 ms, normalized collision energy (NCE) of 35, and dynamic exclusion of 30 s.

#### Data analysis

The MS/MS spectra were searched against the UniProt human reference proteome with the Andromeda 252 search engine integrated into the MaxQuant 253 environment (version 1.5.2.8) using the following settings: oxidized methionine (M), TMT-labeled N-term and lysine, acetylation (protein N-term) and deamidation (asparagine and glutamine) were selected as variable modifications, and carbamidomethyl (C) as fixed modifications; precursor mass tolerance was set to 10 ppm; fragment mass tolerance was set to 0.01 Th. The identifications were filtered using a false-discovery rate (FDR) of 0.01 using a target-decoy approach at the protein and peptide level. Only unique peptides were used for quantification and only proteins with at least two unique peptides were reported. Data analysis was performed using Perseus. Protein levels were median centered and log2 normalized. To identify proteins differentially expressed between groups, a Welch’s t-test was performed.

### ChIP-seq

Around 1×10^7^ 501mel cells per IP were double crosslinked in 25mM EGS (Thermo Fisher Scientific, 21565) and 1% formaldehyde (Sigma Aldrich, Cat#47608). Cells were lysed in ChIP lysis buffer (CLB = 10mM Tris HCl pH8, 100mM NaCl, 1mMEDTA pH8, 0.5mM EGTA pH8, 0.1% Sodium Deoxycholate, 0.5% N-lauroylsarcosine, 1% Triton X-100) and sonicated with Diagenode Bioruptor for 15-30 cycles (30” ON +30” OFF max intensity). H3K27ac IP was conducted using 5ug of Ab/IP H3K27Ac Ab (Active Motif, Cat#91193, RRID:AB_2793797) against an IgG control (Cell Signaling Technology, Cat#3900, RRID:AB_1550038) prebound to ProteinA/G magnetic beads (Pierce Thermo Fisher Scientific, Cat#88803). After overnight IP, beads were washed twice in Wash Buffer I (WBI, 20mM Tris HCl pH7.5, 150mM NaCl, 2mM EDTA pH8, 0.1% SDS, 1% Triton X100), Wash Buffer II (WBII, 10mM Tris HCl pH8, 250mM LiCl, 1mM EDTA, 1% Triton X100, 0.7% Sodium Deoxycholate) and TET Buffer (10mM Trist HCl pH8, 1mM EDTA, 0.2% Tween20) and eluted in Elution buffer (EB, 10mM Tris HCl pH8, 300mM NaCl, 5mM EDTA pH8, 0.5% SDS, 10ug RNAse A, 100ug proteinase K). ChIP elutions were purified using MinElute Reaction Cleanup kit (Qiagen, Cat#28204) followed by library preparation using NEBNext Ultra II DNA Library Prep Kit for Illumina (NEB, Cat# E7600S) following manufacture instructions.

<u>Analysis.</u> H3K27ac and MITF fastqs (MITF fastq obtained from GSE61967) were trimmed, filtered and aligned to hg38 human genome using using Trim Galore (RRID:SCR_011847), FastQC (RRID:SCR_014583) and Bowtie 2 (RRID:SCR_016368) (2.4.1). Sam output files were converted into bam using SAMTOOLS (RRID:SCR_002105) (1.16124) and bigwig files were generated using bamcoverage (deeptools 3.5.6125). H3K27ac and MITF peaks were called using MACS (RRID:SCR_013291) (2.2.9.1126) using paired-end mode. IgG served as controls. During all steps, genomic regions were filtered for blacklisted regions.

### Seahorse Metabolic Analysis

30 to 50 K cells were plated on a XF96 Cell Culture Microplate. The Seahorse MitoStress test protocol was followed using a Seahorse XF96 instrument (Agilent). Concentrations of inhibitors injected were as follows: 2uM oligomycin, 500 nM FCCP and 1 uM antimycin A and rotenone (Cayman Chemical Company, Cat#11342, Cat#15218, Cat #19433, Cat#13995, respectively). After the MitoStress test, cells were fixed in the plated and stained using crystal violet and dissolved with Acetic Acid 15%. Absorbance was measured at 565l. Oxygen consumption rate (OCR) values for every condition were normalized to final confluence.

### Glutamine tracing

For ^13^C5-Gln tracing experiments cells were plated in a six-well plate at 2.0 × 105 cells per well and allowed to attach overnight in DMEM. Cells were treated as indicated in figure legends. Next, cells were washed with PBS twice and cultured for 30 mins in DMEM without phenol (Gibco, Cat#A1443001) supplemented with 4 mM 13C5-Gln and 10% dialyzed serum was added. The tracing medium was removed and washed with PBS. Polar metabolites were collected and derivatized using MOX-TBDMS (Thermo Scientific, Cat#TS-45950, Regis Technologies, Cat#77377-52-7; Cat#18162-48-6, respectively) and analyzed using a labeled amino acid standard as previously described 119–122.

### Animal Studies

NOD/SCID/IL2yR-/- male mice (RRID:IMSR_JAX:005557) at 8-12 weeks were used for *in vivo* studies involving injection of 12-273BM STC. Male athymic nude mice (RRID:IMSR_JAX:002019) were used for in vivo studies involving injection of 10-230BM STC and MDA-231-Brm2. Male mice were used exclusively because injection of these short-term cultures into female mice results in overwhelming ovarian tumor burden that causes mice decline before brain metastasis develops.

All experiments were conducted following protocols approved by the NYU Institutional Animal Care Use Committee (IACUC) (protocol number s16-00051). Animals experiments with YUMM3.2 were conducted in AAALAC approved facilities at the University of Utah. All animal protocols were reviewed and approved prior to experimentation by the Institutional Animal Care and Use Committee (IACUC) at the University of Utah.

#### Metastasis in the YUMM3.2 syngeneic model

C57BL/6J (RRID:MGI:3028467) male and female mice were housed in cages containing up to five animals of the same sex and maintained at room temperature. Mice were fed a combination of Teklad Global 2920X and Teklad 3980X (lnotiv), supplemented with DietGel76A and HydroGel (Clear H2O) post-weaning. One-week post-weaning, mice were transitioned to Teklad Global 2920X.

#### In vivo metastasis assays

100,000 cells suspended in 100 μl of PBS were injected with ultrasound guidance (Visualsonics Vevo 770 Ultrasound Imaging System) into the left ventricle of mice anesthetized with isoflurane. Mice were monitored weekly for metastatic progression by in vivo Bioluminescence imaging (BLI). Upon weight loss > 20% of their maximum weight over the course of the experiment, and/or signs of distress (neurological signs, abnormal locomotion) in any experimental mice, experimental endpoint was established. At experimental endpoint, all mice in all experimental groups were euthanized with ketamine/xylazine (see “Perfusion”).

Generally, 12 mice per group underwent intracardiac injection with cancer cells. Figure legends indicate where group size may vary due to unattended death or failed intracardiac injection.

#### In vivo syngeneic allografts

Seven to nine weeks-old glowing head mice were subcutaneously injected into the dorsal area near the scapula with 5×10^4^ Yumm3.2 BrafV600E Cdkn2a-/- Pten-/- cell expressing Cndp1-HA and Akt1E17K GFP-Luciferase in Matrigel and observed for tumor growth. Tumors were visualized and measured weekly using bioluminescence imaging (BLI), and tumor burden was quantified using bioluminescent photon output values and digital caliper measurements up to 2000 mm^3^. The following formulas was used to calculate tumor volume: (Length X Width^2^)/2.

#### In-vivo BLI

10 minutes prior to imaging, luciferin substrate (150 mg/kg) was administered to mice by intraperitoneal injection. Mice were anesthetized with isoflurane (Covetrus, Cat#11695-6777-2) and imaged by IVIS Illumina instrument (PerkinElmer, (RRID:SCR_025239)) for an automatically-determined duration (1-120 sec). Signal was quantified by measurement of total luminescent flux (p/sec/cm2/sr) in drawn brain and body regions of interest using Living Image software. For the Yumm3.2 syngeneic model organ visualization, at euthanasia, the primary tumor as well as lungs, livers and brains were imaged ex vivo using the IVIS Spectrum to confirm luciferase expression and to detect metastases. Living image software (version 4.5.2) was used to compile images.

#### Perfusion

Mice were anesthetized with a ketamine (100 mg/kg, Covetrus Cat#071069) and xylazine (10 mg/kg, Anased, Cat#NDC59399-110-20) cocktail by intraperitoneal injection. The heart was exposed by gross dissection and an incision was made in the right atrium. Subsequently, 10 mL of PBS followed by 10 mL of 4% PFA was injected into the left ventricle.

#### Continuous shRNA-mediated gene silencing

Starting two days prior to intracardiac injection and continuing through the experimental duration, doxycycline hyclate (200 mg/kg/day) was administered to mice in food pellets.

#### Induction of gene silencing in growing metastases

Protocol for *in vivo* metastasis assay (as described above) was performed. Mice were administered standard chow at the beginning of the experiment. 14-17 days (indicated in figure legend) after intracardiac injection of cancer cells, gene silencing was induced by administration of doxycycline hyclate (200 mg/kg/day) in food pellets.

#### Mouse tissue processing

Organs harvested from PFA-perfused mice were fixed in 10% formalin for 72 hours, followed by several washes PBS and resuspension in 70% ethanol for 24 hours. Prior to embedding, brains were sectioned grossly into thirds coronally and livers were sectioned by lobe. Organs were embedded in paraffin and cut into 5 μM thick sections. The embedded brain thirds were sectioned coronally through the entire length of tissue at an interval of 50 μM. For livers, sections were obtained from two levels 100 μM apart. For brain, sections were obtained from 2 evenly spaced levels of the embedded coronal thirds. In total, this resulted in 6 serial coronal brain section levels for metastatic quantification. All sections were stained with H&E.

### Statistical Analysis

Statistical analyses were performed with Prism 9 (RRID:SCR_002798), specific tests are indicated in figure legends. Unless otherwise stated, values are averages and error bars are +/- standard deviation.

## Resource Availabilit

### Lead contact

Further information and requests for resources and reagents should be directed to and will be fulfilled by the lead contact, Eva Hernando.

### Material availability

Requests for resources and reagents should be directed to and will be fulfilled by the lead contact, Eva Hernando

### Data and code availability

RNA-sequencing/Proteomics/Metabolomics data have been deposited at [TBD] as [Database:TBD] and are publicly available as of the date of publication. Additional dataset utilized are detailed indicated in references, figure legends and Methods.

De-Identified human data is available in Supplementary Tables. They are publicly available as of the date of publication until [TBD].

Original western blots images are available as supplementary information in Document S1. All code and packages are available in this paper’s methods

Any additional information required to reanalyze the data reported in this paper is available from the lead contact upon request. Eva Hernando.

## Acknowledgements

We thank the NYU Grossman School of Medicine core facilities, funded by Cancer Center Support Grant (NCI/NIH; 5P30CA016087), including the Preclinical Imaging Laboratory (Youssef Zaim-Wadghiri, Orlando Aristizabal), the Experimental Pathology Research Laboratory (Director, Dr. Cindy Loomis) for tissue processing and histology, the Proteomics Core (Director, Dr. Beatrix Ueberheide) for mass spectrometry data, Metabolomics Core (Director, Dr. Drew Jones) for metabolic profiling, Genomics Technology Center (Director, Adriana Heguy) for transcriptomics. This work was funded by NIH/NCI U54 CA263001, R21CA286244, 5R01CA277425, the V Foundation, and the Melanoma Research Alliance. I.O was supported by NIH Melanoma SPORE grant NCI P50 CA225450. M.L. was funded by the University Research Priority Program (URPP). A.K was supported by T32CA009161 and T32GM136573. V.O-V was supported by 5F31CA250364. P.B. has been funded by a National Cancer Center fellowship and a Melanoma Research Alliance fellowship. S.L.H. is funded by NIH R01CA121118 and the Melanoma Research Alliance. D.E.B is funded by the NCI Predoctoral to Postdoctoral Fellow Transition Award (F99/K00) K99 CA245822-07. Uta Mackensen designed and performed the illustrations that can be found in Figure S7.

## Authors contributions

A.K., M.A.G-M and E.H conceived and designed the study. M.A.G-M, A.K., M.N., V.O-V., O.K., P.B., A. P.M and E.V.S-dM performed experiments. M.A.G-M., A.K., M.N., A.W., P.B., Y.W., M.I., X.K., S.T. and D.E.B analyzed experimental or genomic data. D.E.B., M.P., R.P., R.J.S., S.L.H., C.T.V., K.V.R. contributed to data interpretation. I.O., M.L, E.dS and B.I. provided patients tissues and/or data. A.K., M.A.G-M., D.E.B. interpreted the data. M.A.G-M, A.K., and E.H. wrote the manuscript. All authors read and approved the manuscript.

## Declaration of interest

The authors declare no competing interests.

---

**Figure S1.**
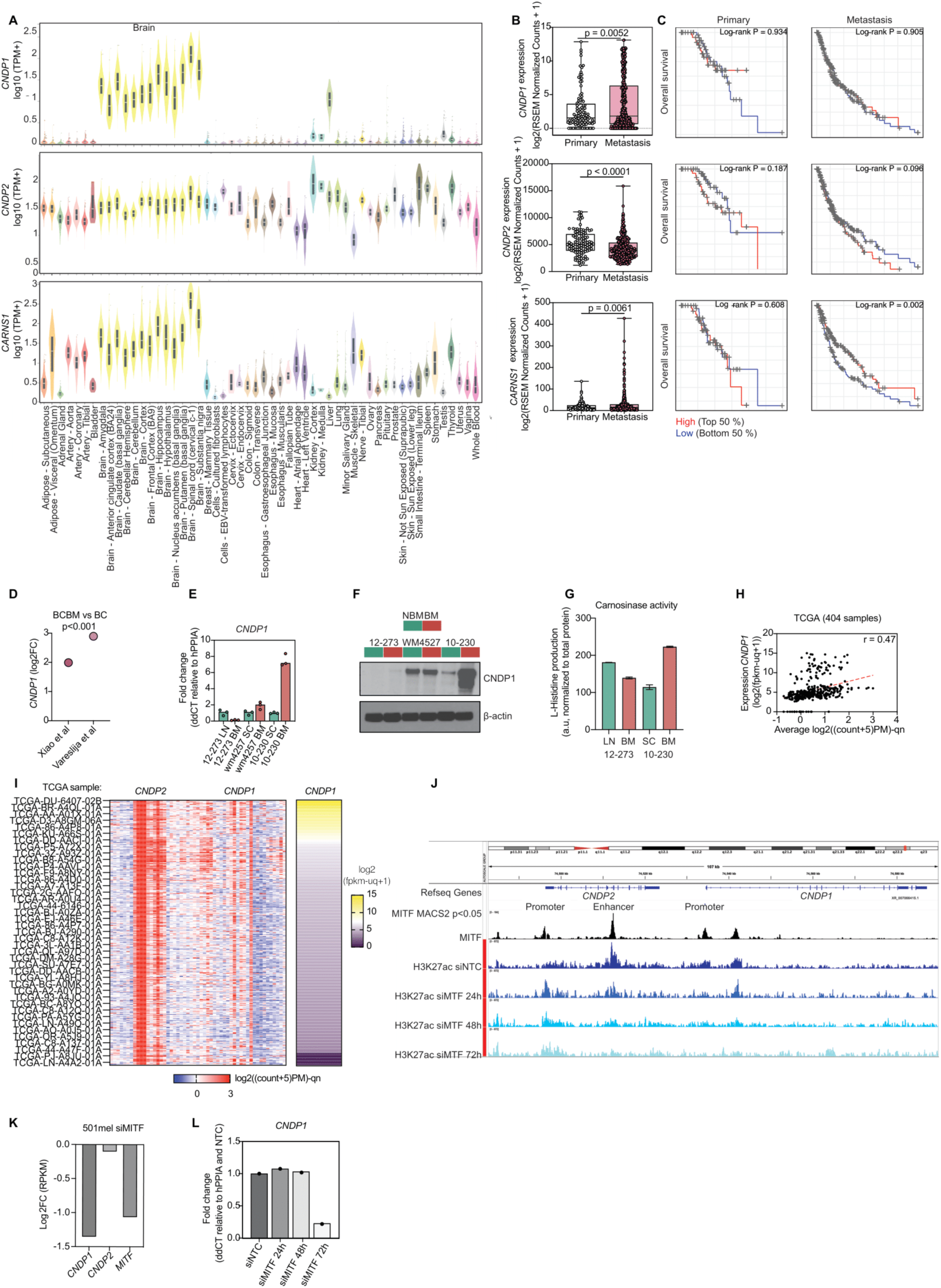
*CNDP1 In silico* analysis in melanoma databases. Related to Figure 1. **(A)** Comparative analysis of log 10 transcription *CNDP1, CNDP2* and *CARNS1* expression levels (TPM+1) in human normal tissues mined from GTEx, the Genotype-Tissue Expression project. **(B)** *CNDP1, CNDP2, CARNS1* (RSEM analysis in normalized counts) expression in primary vs metastatic melanoma human samples from Melanoma dataset Cancer Genome Atlas RNA sequencing (n = 472 total samples from 469 patients. n = 369 metastases and n = 103 primary). P value by unpaired, two-tailed t-test with Welch’s correction. **(C)** Kaplan-Meier curves showing patient Overall Survival stratified by high and low *CNDP1, CNDP2* and *CARNS1* gene expression (RNA-seq) in primary and metastatic tumor on TCGA(113,132,133) Melanoma (SKCM Firehouse Legacy, n = 472) and log-rank test by TIMER2.0(114,131,134). **(D)** 2logFC *CNDP1* expression in breast cancer data bases described in the indicated references (135,136) (adj p value p<0.0001). **(E)** *CNDP1* expression (qRT-PCR) in matched extracranial (NBM) and brain metastasis (BM) derived short term cultures 12-273, WM4527, 10-230. Expression represented as Fold Change relative to housekeeping gene (hPPIA) and to extracranial derived matched short-term culture (n=3). Additional data can be found in Table S1 **(F)** Immunoblot showing CNDP1 expression in patient-matched extracranial (NBM) and brain metastasis (BM) derived short term cultures 12-273BM, WM-4527, 10-230BM. Additional data can be found in Table S1. See also Document S1 for details. **(G)**. L-Histidine liberation assay as carnosinase activity read-out in conditioned media of 10-230 and 12-273 paired matched STC. Additional data can be found in Table S1. Two technical replicates shown. **(H)** *CNDP1* expression significantly correlates with the average epigenetic accessibility around *CNDP1* promoters. In red is shown the linear regression line. R represents the Spearman coefficient. **(I)** Combined heatmap showing the epigenetic accessibility surrounding *CNDP1* and *CNDP2* loci in a subset of TCGA patients (left) and *CNDP1* expression in the corresponding samples (right), (n=404) **(J)** IGV snapshot of MITF and the active chromatin marker H3K27ac genomic occupancy around *CNDP1* and *CNDP2* loci generated in 501mel. MITF binds to *CNDP1* promoter and H3K27ac loss upon MITF silencing at *CNDP1* promoter strongly suggests a direct MITF-dependent CNDP1 regulation. **(K)** Bar plot of the mRNA expression of *CNDP1, CNDP2* and *MITF* upon MITF silencing in 501mel cell. *CNDP1* expression is strongly reduced following MITF depletion compared to *CNDP2* (data extrapolated from Laurette et al., 2015, GSE61967). **(L)** *CNDP1* expression (qRT-PCR) in 501Mel melanoma cells after 24, 48 and 72 h MITF silencing (siMITF). Expression represented as Fold Change relative to housekeeping gene (hPPIA) and non-targeting control silencing (siNTC).

**Figure S2.**
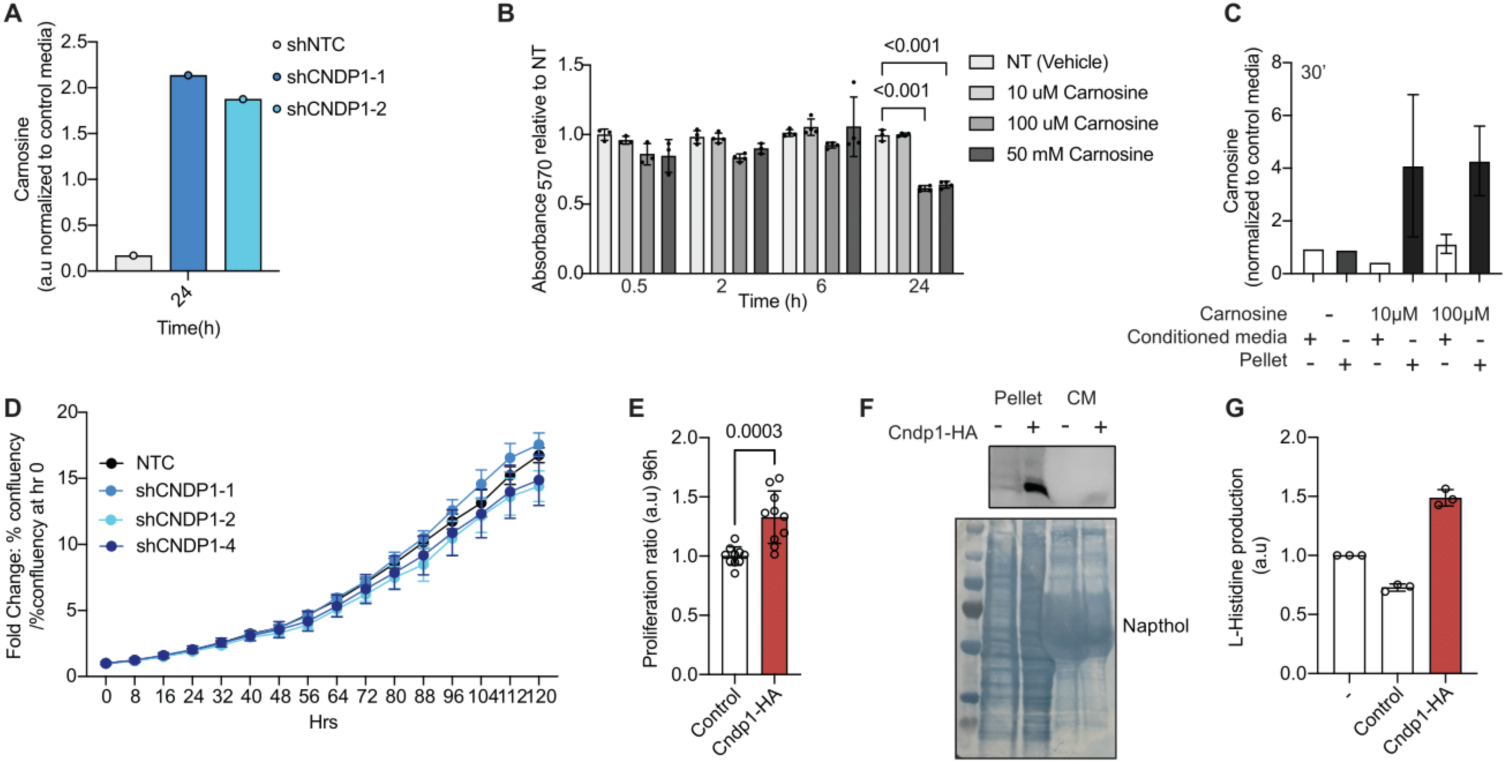
CNDP1 modulation effect in melanoma cells proliferation. Related to Figure 2. **(A)** Carnosine accumulation measured by ELISA represented as normalized values to Carnosine concentration in shNTC conditions (30 mins under culture) in 12-273 transduced with indicated tetON-shRNAs treated with doxycycline for 24h. Mean of two technical replicates shown. **(B)** Proliferation ratio of 10-230 BM cells cultured with 10 μm, 100 μm and 50 mM Carnosine concentrations during 0.5, 2, 6 and 24 h of culture normalized to not treated (NT) of their respective time points (n=3, technical replicates). Confluency obtained by treating the cells with Resazurin and measuring absorbance after 3 hours of treatment at 570 nm. Statistical analysis by one-way ANOVA. **(C)** Carnosine accumulation shown as normalized units to STC media in conditioned media and pellet after 30 minutes of treatment. **(D)** Proliferation ratio of SKMEL-239 cells transfected with indicated shRNAs after 96 h of culture normalized to 0h, indicated as % confluency extracted from Incucyte image analysis. **(E)** Proliferation ratio performed by serial fixing and crystal violet staining of Yumm3.2 BrafV600E/wt Cdkn2a-/- Pten-/- control or ectopically expressing Cndp1-HA after 96h as indicated. **(F)** Immunoblot showing Cndp1-HA expression in conditioned media and pellets of Yumm3.2 BrafV600E/wt Cdkn2a-/- Pten-/- control or ectopically expressing Cndp1-HA. See also Document S1 for details. **(G)** L-Histidine liberation assay as carnosinase activity read-out in pellets of Yumm3.2 BrafV600E/wt Cdkn2a-/- Pten-/- control or ectopically expressing Cndp1-HA. Three technical replicates shown. One representative technical replicate shown for all the above assays.

**Figure S3.**
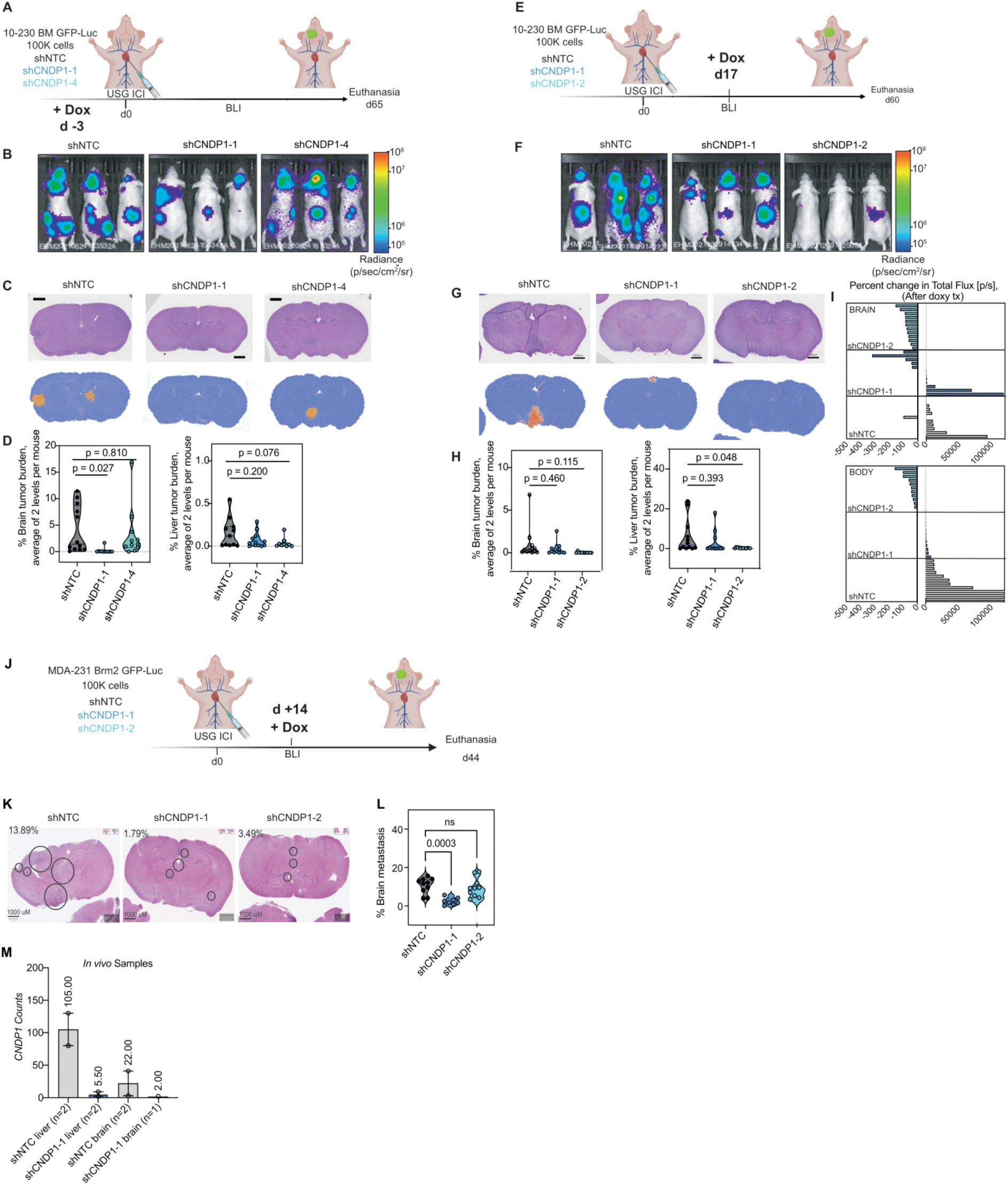
CNDP1 Silencing Decreases *In Vivo* Melanoma Metastasis. Related to Figure 3. **(A)** Schematic representation of the *in vivo* xenograft constitutive approach experiment (USG ICI). **(B)** Representative images of whole-body *in vivo* BLI of athymic/nude mice injected with 10-230 BM cells transduced with indicated tetON-shRNA 52 days post-injection (n = 12 mice per condition, BLI represented by radiance: p/sec/cm^2^/sr). **(C)** Representative H&E and MetFinder brain metastasis sections of nude mice injected with indicated (n = 12 mice per condition, duplicated sections and levels analyzed per organ/mice, Scale bar 1000 μm and 2000 μm for brain and liver, respectively). **(D)** Tumor burden in brain, leptomeningeal and liver quantified as the percent of area occupied by the tumor of the total organ area and averaged of 2 levels of sectioning per organ obtained by performing MetFinder analysis. Statistical analysis by one-way ANOVA. **(E)** Schematic representation of the *in vivo* xenograft therapeutic approach experiment (USG ICI: Ultra Sound Guided Intra-Cardiac Injection). **(F)** Representative images of whole-body *in vivo* BLI of nude mice injected with 10-230 BM cells transduced with indicated tetON-shRNA day 59 post-injection (n = 12 mice per condition, BLI expressed as radiance: p/sec/cm^2^/sr). **(G)** Representative H&E and MetFinder brain and liver metastasis sections of NSG mice injected with indicated (n = 12 mice per condition, duplicated sections and levels analyzed per organ/mice, Scale bar 1000 μm and 2000 μm for brain and liver, respectively). **(H)** Tumor burden in brain, leptomeningeal and liver quantified as the percent of area occupied by the tumor in the total organ area and averaged of 2 levels of sectioning per organ. Obtained by performing MetFinder analysis. Statistical analysis by one-way ANOVA. **(I)** Percent change in IVIS signal in the brain (top) and whole body (bottom) from days 12 to 36 of nude mice injected with 10-230 BM cells transduced with indicated tetON-shRNA. One bar = one mouse, n = 10 mice/group. **(J)** Schematic representation of the *in vivo* xenograft therapeutic approach experiment (USG ICI: Ultra Sound Guided Intra-Cardiac Injection). **(K)** Representative H&E images and corresponding quantitation of metastasis burden in brain sections of NSG mice injected with MDA-231 BrM2 cells transduced with indicated tetON-shRNA obtained by manual analysis. Not burden in lung, liver or kidneys detected. **(L)** Tumor burden in brain quantified as the percent of area occupied by tumor and average of two levels of sectioning per organ (100 µM separation) obtained by manual analysis (n=10, 8, and 12 mice per group, two sections and two levels analyzed per organ/mice). Statistical test: one-way ANOVA. Scale bars represent 1000 µm. Not burden in lung, liver or kidneys detected. **(M)** *CNDP1* expression (counts) in RNA sequencing of 12-273 BM GFP positive cells harvested from 4 mice livers and 3 mice brains of the indicated conditions after USG ICI and 7 days of doxycycline treatment.

**Figure S4.**
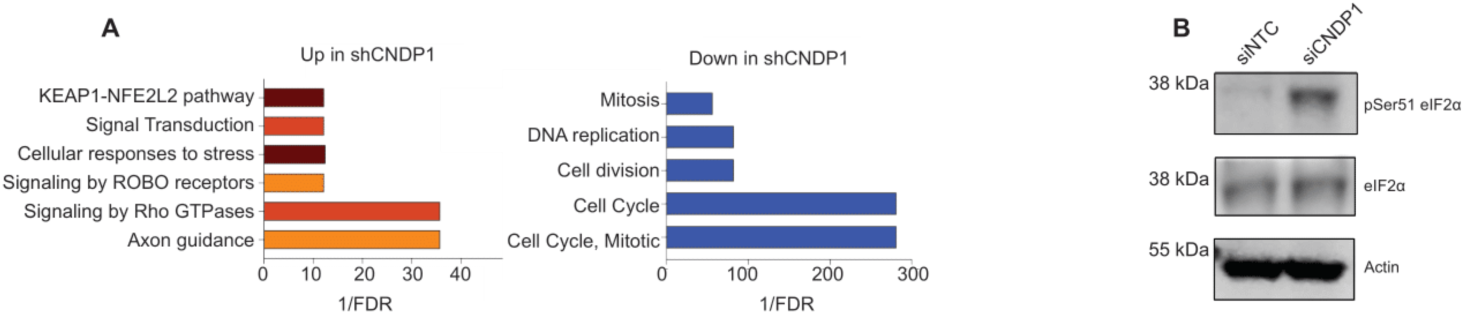
Proteomics changes upon CNDP1 silencing in melanoma cells. Related to Figure 4. **(A)** Representation of DAVID functional annotation algorithm of upregulated (right) and downregulated (left) differentially expressed proteins identified by unpaired, two-tailed t-test comparing 2 groups: shCNDP1-1 and shCNDP1-2 versus shNTC. Represented by 1/FDR as indicated. See Table S10. **(B)** Representative immunoblots of phophoSerine51 eIF2α and total eIF2α, housekeeping (HK) protein b-actin in lysates of 10-230 BM cells transfected with siRNA NTC or CNDP1 as indicated. Results for HRI and translation related proteins expression from samples shown in Figure 6K and Figure 5B, respectively. Total and phospho-proteins were blotted in different membranes. Representative HK shown. See Document S1 for details.

**Figure S5.**
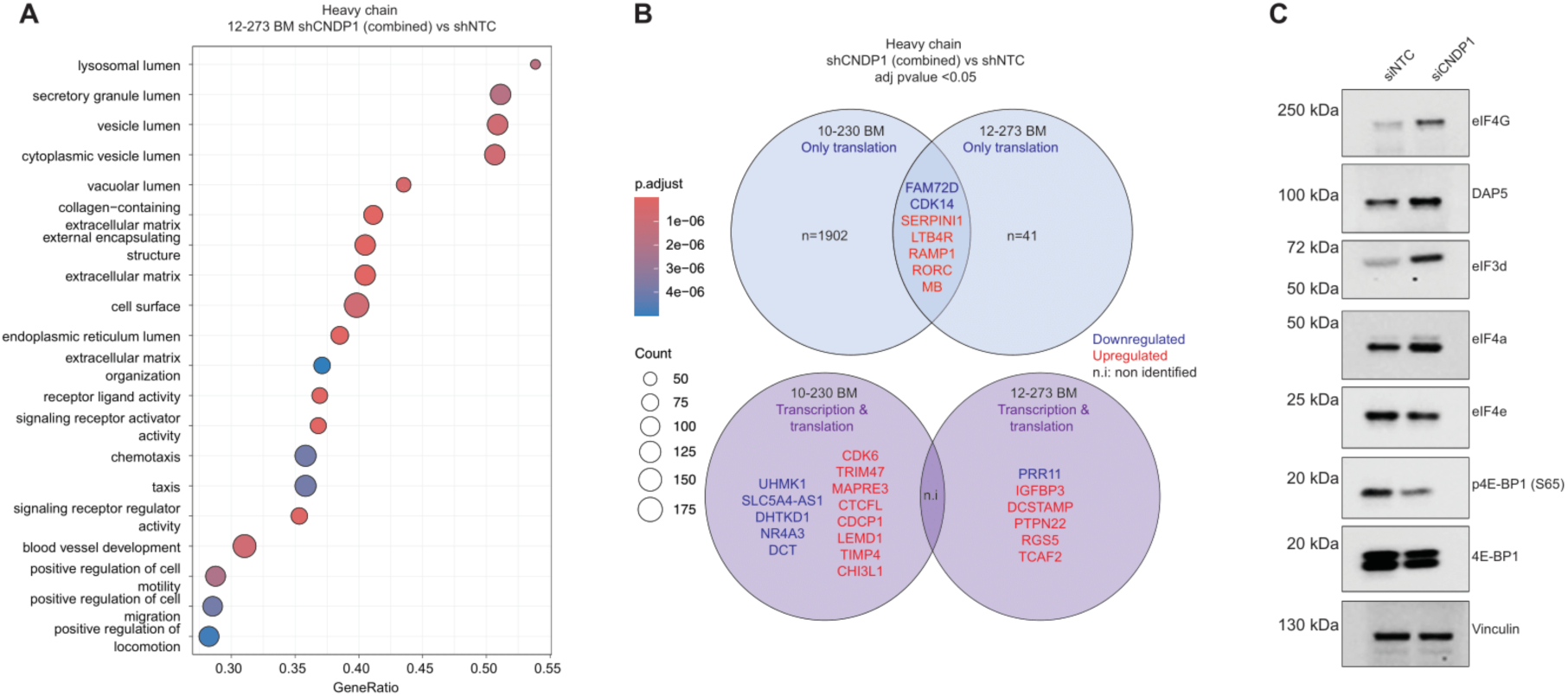
Extended characterization of translation phenotype of CNDP1 KD melanoma cells. Related to Figure 5. **(A)** Integrated Gene Enrichment analysis of 12-273 BM heavy chain both shCNDP1 vs shNTC RNA sequencing represented by Normalized Enrichment Score using Reactome Gene Ontology Analysis (RNA-seq: adjust p value < 0.05). **(B)** Venn diagram including common transcripts changing as indicated at translation level (top) and transcription and translation level (bottom) in both 10-230 and 12-273 BM shCNDP1 (combined) vs shNTC polysome heavy chain (adjusted p value <0.05, n.i -non-identified). See Table S11. **(C)** Representative immunoblot of eIFs proteins and phophoSerine51 4E-BP1, 4E-BP1 and housekeeping (HK) protein Vinculin in lysates of 12-273 BM cells transfected with siRNA NTC or CNDP1 as indicated. See Document S1 for details.

**Figure S6.**
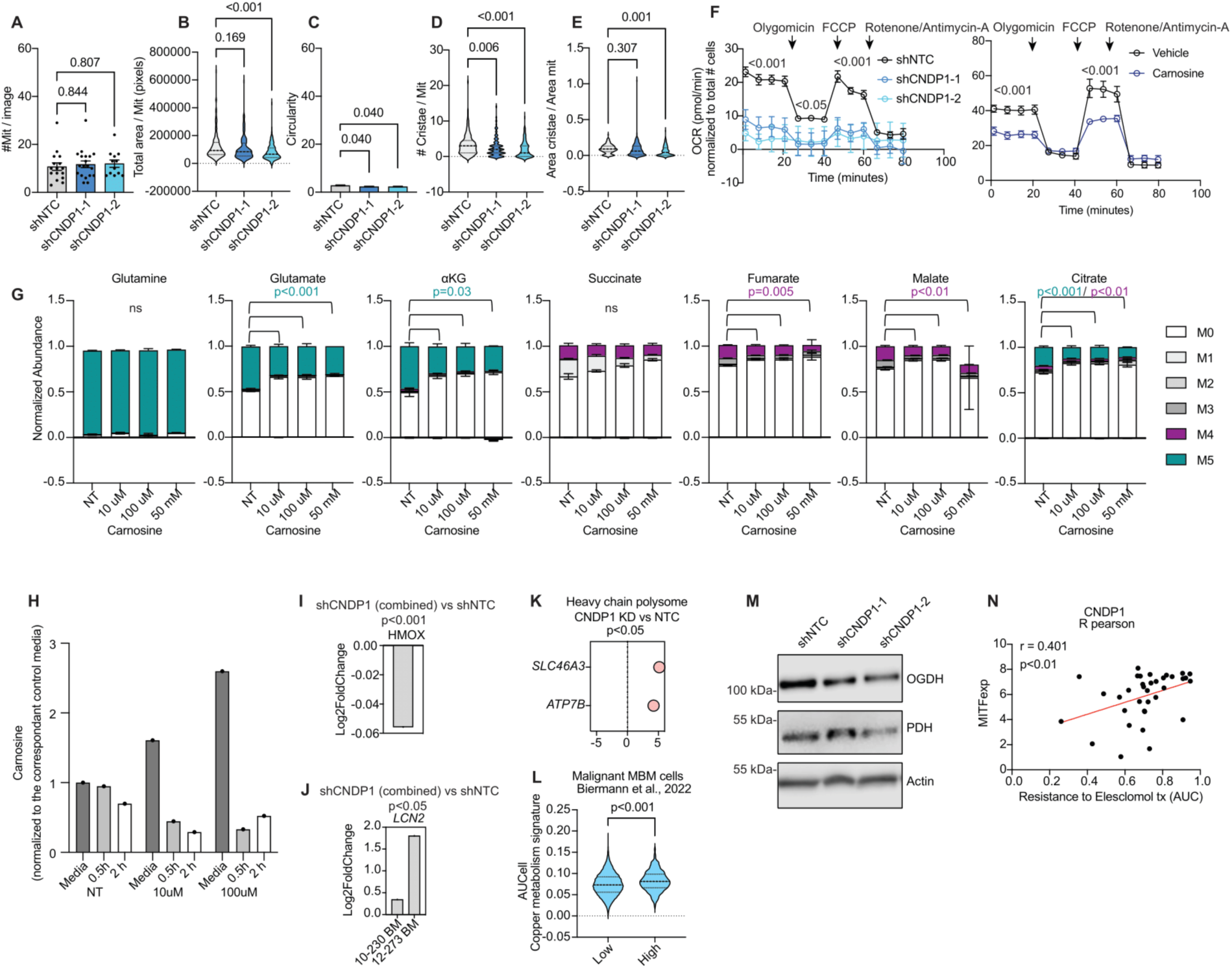
Extended characterization of mitochondrial phenotype of CNDP1 KD melanoma cells. Related to Figure 6. **(A-E)** Quantification of total mitochondria per cell **(A)**, mitochondria area **(B)**, circularity **(C)**, cristae number per mitochondria **(D)** and cristae area per mitochondria area **(E)** in 12-273 BM. Statistical analysis by one-way ANOVA. **(F)** Seahorse MitoStress analysis of OCR in 10-230 BM transduced with indicated tetON-shRNA and MDA-231 Brm2 treated as indicated. Statistical analysis by one-way ANOVA. Representative replicate shown (n=3, p<0.05 or indicated). **(G)** Normalized distributions of glutamine, glutamate, and TCA cycle metabolites, including α-ketoglutarate (αKG), succinate, fumarate, malate and citrate, in 10-230 BM cells treated with 10 10 μm, 100 μm and 50 mM Carnosine for 30 minutes are shown. (n = 3); data are shown as the mean ± SD. Statistical analysis by one-way ANOVA. P value indicated. Cells were cultured with [U-^13^C5] glutamine for 6 h before metabolite extraction and gas chromatography-mass spectrometry (GC-MS) analyses. **(H)** Carnosine accumulation measured by ELISA represented as normalized values to carnosine concentration in dialyzed media supplemented with indicated concentrations in 10-230 BM cells cultured with 10 μm, 100 μm and 50 mM Carnosine concentrations during 0.5, 2 h. Mean of two technical replicates shown. Results for 0.5 h also shown in Figure S2C. **(I)** 2logFold Change of HMOX protein abundance measured by proteomics (MS/MS) in 10-230 BM cells transduced with indicated tetON-shRNAs. Differentially expressed proteins identified by unpaired, two-tailed t-test comparing 2 groups: shCNDP1-1 and shCNDP1-2 versus shNTC. **(J)** 2logFoldChange of *LCN2* transcript expression of both shCNDP1 tetON-shRNA vs shNTC in 10-230 BM and 12-230 BM cells. RNA sequencing data. p value < 0.05. **(K)** 2logFold Change of *ATP7B* and *SLC46A3* transcript of Heavy Chain RNA sequencing of 10-230 BM shCNDP1 vs shNTC. p value < 0.05. **(L)** AUCell (Area Under the Curve) measurement of copper homeostasis gene signature (WikiPaths, WP3286) in *CNDP1* high expression versus *CNDP1* low expression in MBM cells from Biermann et al., 2022 data mining. Statistical analysis by one-way ANOVA, p<0.001. **(M)** Representative immunoblots of OGDH, PDH and housekeeping (HK) protein b-actin in lysates of 10-230 BM cells transfected with indicated sh-RNAs. See document S1 for details. **(N)** R Pearson correlation between melanoma cell lines classified by their MITF expression and resistance to Elesclomol treatment expressed as AUC of cell viability after treatment. Data mining from Depmap.

**Figure S7.**
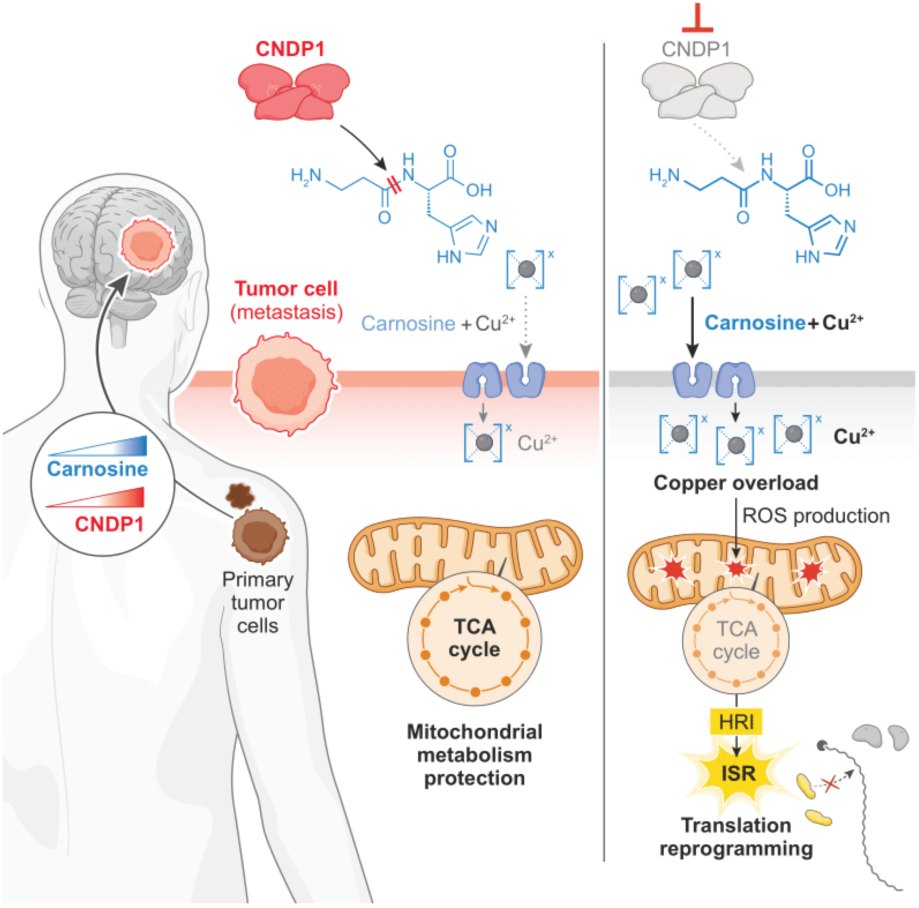
Graphical representation of CNDP1 role in brain metastasis.

## Notes

### Competing Interest Statement

The authors have declared no competing interest.

### Summary of Updates

Figure 1G - error in duplicate images. Update version with the right images

